# Seizures and tauopathy following neurotrauma are mediated by prion protein and metabotropic glutamate receptor 5

**DOI:** 10.64898/2026.09.21.753012

**Authors:** Laszlo F. Locskai, Janelle Nordin, Taylor Gill, Melissa J. Kinley, Hadeel Alyenbaawi, W. Ted Allison

## Abstract

Traumatic brain injury (TBI) is one of the world’s leading causes of death and disability and a major risk factor for dementias. The primary dementia associated with TBI is chronic traumatic encephalopathy (CTE), a neurodegenerative disease classified as a tauopathy, in which toxic tau molecules lead to disease pathologies and degeneration. The processes that lead to tauopathy and subsequent dementia after TBI remain unclear. Here, we built upon the finding that seizures after TBI may be a mechanism leading to tauopathy, by dissecting the functions of the metabotropic glutamate receptor 5 – cellular prion protein (mGluR5-PrP^C^) pathway. We delivered TBI to larval in a blast paradigm, and quantified aggregation of Tau via a genetically-encoded Tau-GFP fusion reporter. Zebrafish larvae lacking *prp2* (homolog of mammalian cellular Prion Protein, PrP^C^) displayed a 168% increase in post-traumatic seizures activity after TBI. An mGluR5 agonist (CHPG) reduced post-traumatic seizures, whereas an mGluR5 antagonist (MPEP) increased post-traumatic seizures. Moreover, agonizing mGluR5 reduced tau aggregation and antagonizing mGluR5 increased tau burden. Larvae seizing from convulsants, rather than TBI, were treated with CHPG/MPEP and provided a similar pattern of outcomes, suggesting seizures may be a factor needed for mGluR5 activity to influence tau aggregation. The PrP^C^-mGluR5 pathway is proposed as one candidate pathomechanism linking TBI to subsequent seizures and tauopathy, and thus it warrants investigation as a target for prophylactic interventions.

## Introduction

Traumatic brain injuries (TBI) impact people from many walks of life including children, professional athletes, domestic abuse victims, military personnel, and first responders^1–4^. TBI is defined as a physical insult to the brain through direct impact, rapid movements, or blast waves; and these range from mild to severe forms of injuries^5^. TBI is the number one cause of death and disability under the age of forty-five^6,7^, impacting an estimated 69 million people per year worldwide^8^. TBI is also a major risk factor for tragic neurodegenerative diseases and dementia, such as chronic traumatic encephalopathy (CTE) and Alzheimer’s disease (AD), which are limited in treatment options and are poorly understood^9–12^. Both diseases are classified as tauopathies which involve the accumulation of pathological tau protein, axonal degradation, and neuronal loss^13–15^. The pathomechanisms linking TBI to tauopathy have been enigmatic, but recent research has revealed that neuronal hyperactivity^16–20^ and seizures^15,21^ may be an accessible link that drives tauopathy after TBI.

Multiple studies exist displaying a connection between neural activity and seizures with an increase in tau pathology. Increased neuronal activity and excitotoxicity has been shown to increase the release of tau protein^16–18^ and abundance of pathological tau^22,23^.The increased release of tau protein mediated by neural activity has been shown to contribute to increased tau pathology^19^. Increased synchronous brain activity has also been associated with increased tauopathy and the brain regions with higher neural activity appear to contain higher loads of tauopathy^20^. Multiple seizure models have also been shown to increase tauopathy^20,21,24^, with evidence to suggest that blocking seizures can prevent tauopathy^21^. Furthermore, epilepsy patients can have tau pathology that is associated with cognitive decline^25–27^, with some suggesting that seizure-mediated cell death releases pathogenic soluble tau^28^. Interestingly, tau has also been shown to increase seizures in animal models, where removal of tau or hyperphosphorylated tau reduces seizure incidents^29,30^. Taken together, this data suggests that increased neural activity/seizures increase tauopathy, while tauopathy simultaneously increases seizures creating a vicious cycle contributing to the cognitive decline seen in multiple diseases. Understanding the mechanisms of how seizures contribute to the development of tauopathy could provide an avenue for early prophylactic prevention of tau-based dementias and diseases.

One pathway of interest, that is strikingly linked to both seizures and tauopathy, is the cellular prion protein (PrP^C^) and metabotropic glutamate receptor 5 (mGluR5) pathway. PrP^C^ is a membrane-anchored glycoprotein most known for its involvement in neurodegenerative prion diseases^31^. mGluR5 is a group I metabotropic glutamate receptor which has multiple signaling cascades through G_q_ /G_11_ G proteins^32^ and has been shown to interact with PrP^C^ at the membrane, causing specific signal transduction^33,34^. PrP^C^ has two functional involvements important to seizures and subsequent tauopathy. Firstly, the loss of function of PrP^C^ and mGluR5 are key components of a rheostat that controls seizure susceptibility in animal models^35–37^. Secondly, PrP^C^ acts as a high-affinity receptor that binds Aβ oligomers; this leads to disease pathology, sleep disruption and increased tau hyperphosphorylation, which are mediated through mGluR5^38–40^. Preventing the toxic signaling mediated by the binding of Aβ oligomers to PrP^C^ has been shown to reduce tauopathy in AD mice models, making this pathway an attractive target for therapeutic development^41^.

In this study, we examined if the disruption of PrP^C^ and/or mGluR5 contributes to TBI pathology and tauopathy using genetic and pharmaceutical interventions. Our goals were to learn how TBI-mediated pathology leads to seizures and to dissect the pathomechanism of TBI-induced seizures that cause tauopathy. We hypothesized that the PrP^C^-mGluR5 pathway functions as a connective mechanism between TBI, seizures, and tauopathy. To model TBI, we sealed larval zebrafish in a syringe and dropped a weight onto the plunger of the syringe. This simple yet elegant TBI method causes blast wave TBI injuries, leading to pathological markers like neuronal cell death, decreased blood flow, seizures and the development of Tau aggregates^21^. To assess tauopathy in larval zebrafish, we used our Tau biosensor larvae, which express a truncated human Tau-4R domain linked to GFP. Our tauopathy model has been previously validated and can detect Tau aggregation after the injection of pathological Tau variants, or after TBI, through the quantification of GFP-positive aggregates^21^.

Our findings suggest that inhibiting mGluR5-PrP^C^ is protective in the presence of Aβ oligomers. Yet, PrP^C^ and mGluR5 both impact the severity of post traumatic seizures and loss of PrP^C^ and/or mGluR5 inhibition increase tau aggregation after TBI. Our results are broadly consistent with the expected roles of mGluR5 defined in AD research and our Aβ tauopathy model, and they are consistent with the functions of mGluR5 related to TBI and seizures. In AD pathology, Aβ oligomers trigger toxic events through PrP^C^-mGluR5 mediated Fyn kinase signalling, indicating that mGluR5 inhibition is protective^38,41^. In TBI research, activating mGluR5 via the agonist CHPG has been shown to be neuroprotective through a separate pathway involving PI3K/Akt signalling^42^. Our results further support the neuroprotective role of mGluR5-PI3K/Akt signalling in a novel way by assessing its role in TBI mediated tauopathy, while also highlighting the need to consider different pathophysiological events when developing potential therapeutics. This connection between the PrP^C^-mGluR5 pathway, seizures, and tauopathy reveals a possible mechanistic target for treating post traumatic seizures and subsequent dementias.

## Results

### Inhibiting mGluR5 signalling reduces Aβ_1-42_ oligomer mediated tauopathy

mGluR5 has been shown to interact with Aβ oligomers through PrP^C^ which contacts mGluR5 at the cell membrane. Aβ oligomers can bind PrP^C^ in a ligand-receptor capacity, resulting in toxic signaling events through mGluR5 which lead to AD pathologies such as increased tauopathy^38,39,41^. Inhibiting toxic mGluR5 signaling and preventing the Aβ oligomer-PrP^C^ interaction have both been shown to stop Aβ oligomer mediated pathology^38,41^. We first determined if Aβ oligomers mediate tauopathy in our larval zebrafish model. To assess this, we injected Aβ_1-42_ oligomers into the hindbrain ventricle of zebrafish larvae and quantified the number of Tau aggregates that formed using zebrafish GFP linked to the human Tau4R domain. Zebrafish larvae injected with Aβ_1-42_ oligomers had a statistically significant increase in GFP+ Tau aggregates in the spine compared to un-injected larvae (Fig. 1A and 1C, p<0.0001; Supplemental Figure S1). Aβ <u>monomers</u> should not be able to increase tauopathy through PrP^C^-mGluR5 signalling, so we next injected Aβ_1-42_ monomers into zebrafish larvae to test the specificity of the Aβ oligomer pathology. Larvae injected with Aβ_1-42_ monomers did not show an increase in GFP+ Tau aggregates, with levels comparable to un-injected larvae (Fig. 1A and Supplemental Figure S1A).

**Figure 1.**
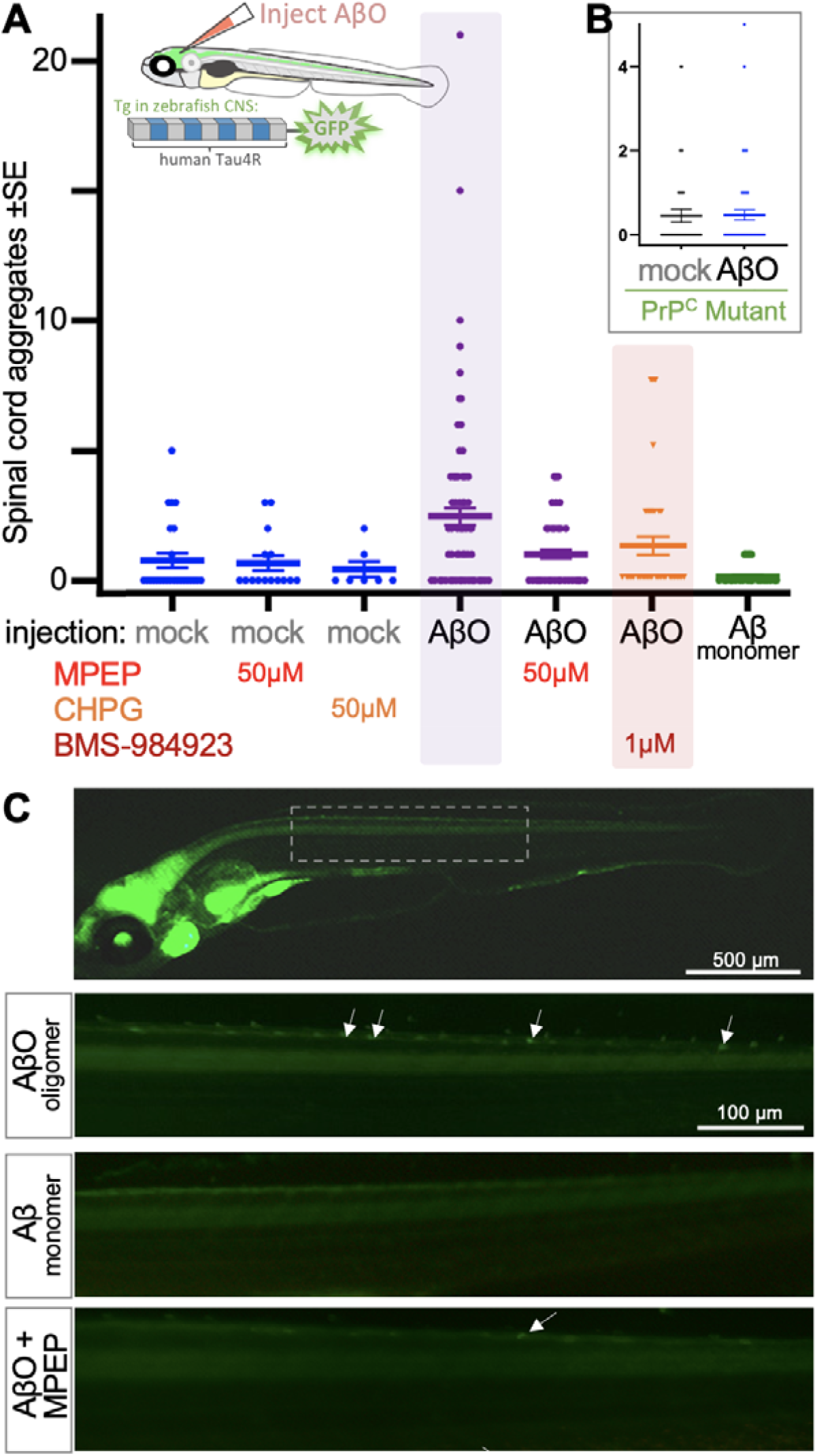
Inhibiting metabotropic glutamate receptor 5 (mGluR5) signalling reduces tau aggregation induced by amyloid beta_1-42_ oligomers. Schematic of “tau biosensor” larval zebrafish with at fusion of human Tau repeat region to GFP expressed in CNS neurons. GFP+ tau puncta are induced by injection with human amyloid beta Aβ_1-42_ oligomers (AβO). Injection at 3 days post-fertilization and Tau quantification 4 days later. **A**. GFP+ tau puncta (each data point represents abundance in an individual animal) are induced by Aβ_1-42_ oligomers (p<0.001) but not monomers. mGluR5 manipulations with antagonist 2-Chloro-5-hydroxyphenylglycine (CHPG, antagonist) reduced tau aggregates to baseline levels, whereas agonist 2-Methyl-6-(phenylethynyl)pyridine (MPEP) had no discernable effect. Solid lines represent the mean ± standard error. **See Supplemental Figure S1 for statistical comparisons and further treatment combinations**. BMS-984923, an mGluR5 silent allosteric modulator that blocks mGluR5’s interaction with cellular prion protein (PrP^C^) complexed to AβO, reduced the impact of AβO injections on tau aggregation. **B**. The impact of AβO injections on tau aggregation depend upon the presence of PrP^C^. **C**. Exemplar images of larval zebrafish with Tau4R-GFP biosensor in CNS neurons (anterior to the left). Top panel denotes inset displayed in lower panels.

After establishing that Aβ oligomer-mediated tau aggregation is observed in larval zebrafish, we next wanted to test whether the toxic Aβ oligomer-mediated signaling through PrP^C^-mGluR5 is conserved in larval zebrafish, and if it is required for the increased tauopathy detected. To establish if toxic PrP^C^-mGluR5 signaling was present, we treated Aβ_1-42_ oligomer injected larvae with the mGluR5 antagonist MPEP, which has been shown to prevent Aβ oligomer mediated pathology^38^. Aβ_1-42_ oligomer-injected larvae with decreased mGluR5 signalling showed a 59% decrease in GFP+ Tau aggregates compared to non-MPEP treated Aβ_1-42_ oligomer-injected larvae and were not statistically higher than un-injected controls (Fig.1A and C, Supplemental Figure S1A). We additionally tested the mGluR5 agonist CHPG on Aβ_1-42_ injected larvae but found no significant changes in the amount of GFP+ Tau aggregates when mGluR5 signalling was increased via CHPG (Supplemental Figure S1A). These results show that Aβ oligomer mediated pathology through mGluR5 signalling is conserved in larval zebrafish.

We next injected *prp2*^−/−^ zebrafish larvae (harboring disrupted PrP^C^) with Aβ_1-42_ oligomers to further test if the increased tauopathy caused by Aβ_1-42_ oligomers occurs through the PrP^C^-mGluR5 pathway (Supplemental Figure S1C). Quantification of Tau4R-GFP suggested that Aβ_1-42_ oligomer-injected *prp2*^−/−^ larvae display no increased tauopathy compared to the un-injected group (Fig. 1B). In addition, we applied the silent allosteric modulator BMS-984923 (BMS) on wild-type fish as a second method to disrupt the PrP^C^-mGluR5 interaction. BMS has been previously shown to alter the conformation of mGluR5 preventing PrP^C^ binding without altering endogenous mGluR5 signalling^41^. BMS treatment prevented Aβ oligomer-mediated tauopathy in a dose-dependent manner (Fig.1A, Supplemental Figure S1B). These results are consistent that Aβ oligomers mediating tauopathy through PrP^C^ in zebrafish. Taken together with the data above, the PrP^C^-mGluR5 pathway appears to be deeply conserved in zebrafish larvae, supporting this as an appropriate preclinical model to investigate the role of the PrP^C^-mGluR5 pathway following insults that promote tauopathy.

### mGluR5 signalling reduces GFP+ Tau aggregates in transgenic zebrafish larvae after TBI

After establishing that Aβ oligomers promote tauopathy through the PrP^C^-mGluR5 pathway in our larval zebrafish model, we next wanted to determine if inhibiting mGluR5 signaling is protective against TBI-induced tauopathy. To do so, we subjected our Tau biosensor larvae to TBI and quantified the number of GFP+ Tau aggregates after pharmacologically increasing or decreasing mGluR5 signalling (schematized in Figure 2A-C). Decreasing mGluR5 signalling after TBI caused a dose dependent increase in GFP+ Tau aggregates (Fig. 2D,E). The largest doses of MPEP (50µM and 100µM) had a 2-fold increase versus larvae with TBI but no drug treatment (p<0.05 TBI + 50uM vs TBI, Fig. 2E). Interestingly, decreasing mGluR5 signalling did not have any significant effects on GFP+ puncta without TBI (Fig. 2E, grey). In summary, animals with altered mGluR5 signalling were hypersensitive to TBI insomuch that they subsequently developed a greater abundance of Tau aggregates.

**Figure 2.**
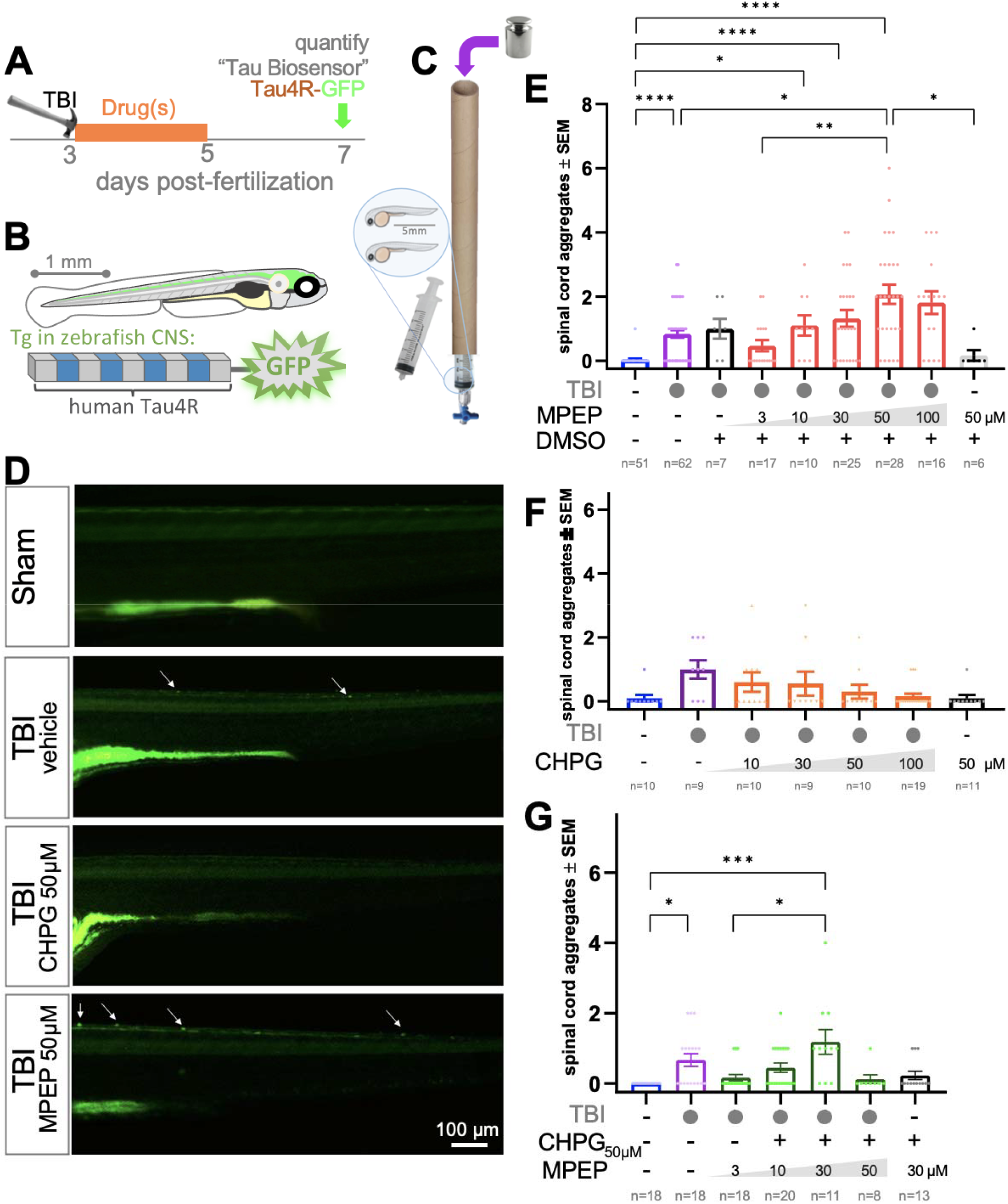
Increased metabotropic glutamate receptor 5 (mGluR5) signaling decreases Tau aggregation in Zebrafish after traumatic brain injury (TBI). A. Larvae were subjected to TBI at 3dpf and treated with drugs for 38 hours. At 7dpf the larvae were quantified via fluorescence microscopy by blinded observers. Each data point represents an individual animal. **B**. Transgenic larvae express human tau repeat region (Tau4R) fuse to GFP in their CNS neurons. **C**. The blast TBI paradigm seals zebrafish larvae in a syringe and delivers a controlled weight drop, creating a pressure wave through the whole body. **D**. Exemplar images of tau aggregation (arrows) in the CNS, most easily visualized in the spinal cord. **E**. The number of spinal GFP+ tau puncta in increases significantly in a dose-dependent manner when mGluR5 signalling is decreased after TBI. 3dpf larvae were treated with 2-Methyl-6-(phenylethynyl)pyridine (MPEP) for 38hrs post-TBI. **F**. The number of spinal GFP+ tau aggregates decreases significantly in a dose-dependent manner when mGluR5 signalling is increased after TBI. 3dpf larvae were treated with 2-Chloro-5-hydroxyphenylglycine (CHPG) for 38hrs post TBI. **G**. MPEP counteracts CHPG’s ability to decrease tau aggregation after TBI in a dose dependent manner, supporting that both drugs are acting through mGluR5. n = number of animals, each experimental group was replicated at least twice. Bars represent the mean ± standard error. Data was analyzed using a Kruskal-Wallis test with Dunn’s multiple comparison. Symbols indicate statistical significance: ^*^<0.05, ^***^<0.001, ^****^<0.0001. Exact p-values and statistical details are listed in Supplemental Tables.

Increasing mGluR5 signalling after TBI had directly opposite results displaying a dose dependent decrease in GFP+ Tau puncta (Fig. 2D and F), reaching levels similar to the no TBI control at 100µM CHPG (Fig. 2F). CHPG treatments did not statistically alter the outcomes. To challenge the specificity of our mGluR5 pharmacology, we used a concerted application of MPEP and CHPG: We held CHPG at a constant dose of 50µM while increasing the concentration of MPEP. CHPG reduced TBI-induced tauopathy as above, and this was reversed by co-application of MPEP in a dose-dependent fashion (Fig. 2G, compare to 2E). These results support the specificity of the applied pharmacology, and thus mGluR5’s role in promoting Tau aggregation following TBI. Overall, the data in Figure 2 support that mGluR5 activity after TBI is protective against Tau aggregate formation, rather than contributing to toxicity as seen in Aβ oligomer mediated pathology.

### mGluR5 signalling is neuroprotective after traumatic brain injury

To further examine if mGluR5 signaling reduced other markers of tauopathy and neurodegeneration, we measured neuronal cell death after TBI by detecting anti-active-caspase-3 levels in zebrafish with increased or decreased mGluR5 activity. We predicted that increased mGluR5 signaling via CHPG would be neuroprotective based on our results that mGluR5 signaling reduces Tau puncta (Fig. 2G) and previous reports of CHPG being neuroprotective after TBI^42–45^. Indeed, increasing mGluR5 signalling after TBI decreased CNS cell death to levels similar to larvae without TBI which was significantly lower (p<0.01) than the CNS cell death in non-treated larvae after TBI (Fig. 3). Likewise, inhibiting mGluR5 signaling after TBI resulted in a 2-fold increase in CNS cell death compared to untreated TBI larvae (p<0.0001, Fig. 3). The observation of mGluR5 signaling after TBI reducing CNS cell death further strengthens the above data that mGluR5 signaling after TBI is neuroprotective and reduces markers of tauopathy.

**Figure 3.**
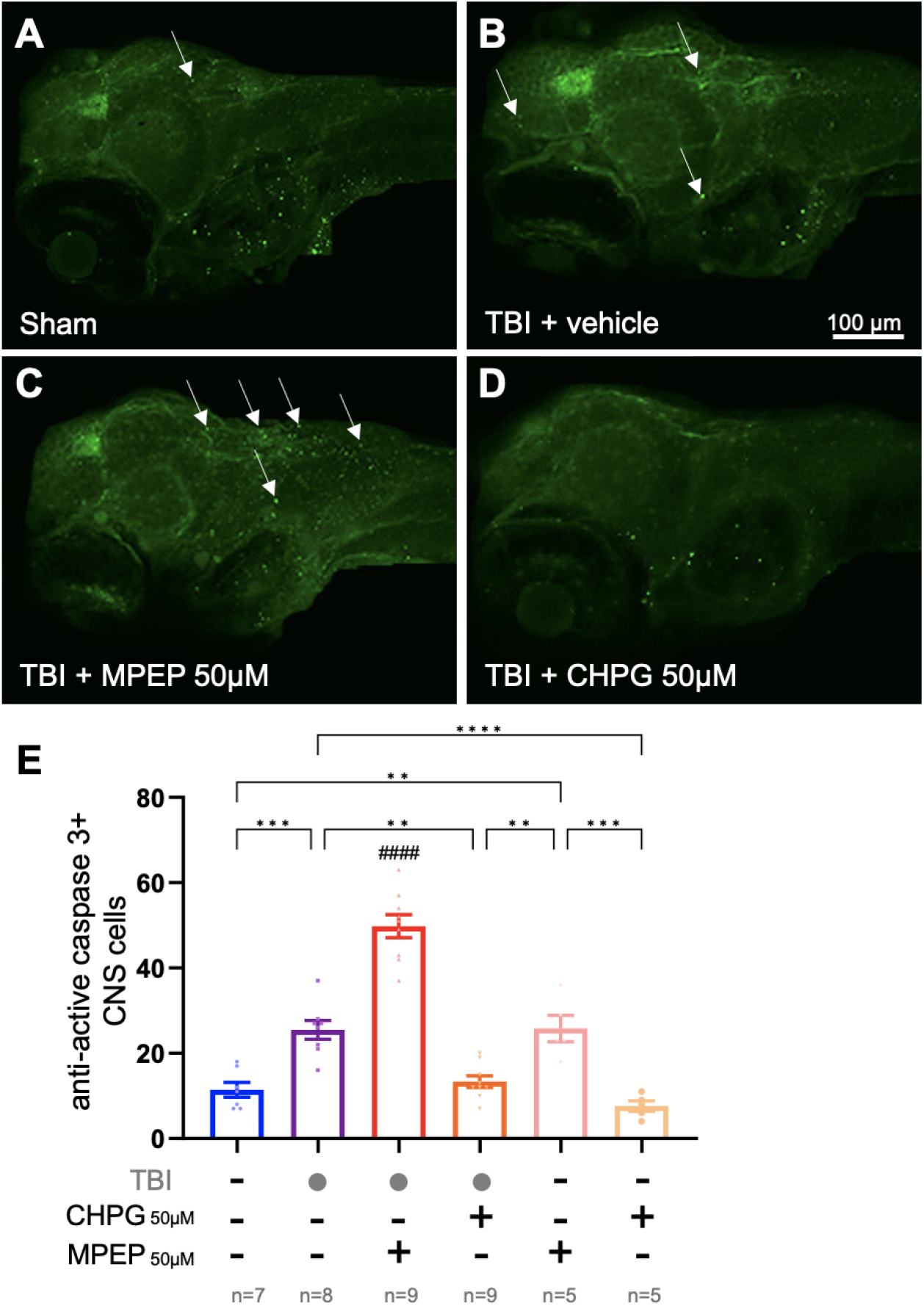
Metabotropic glutamate receptor 5 (mGluR5) signalling is neuroprotective after traumatic brain injury (TBI). Larvae were subjected to TBI at 3dpf and treated with drugs for 38 hours. At 7dpf the larvae were fixed and stained with anti-active-caspase-3 antibody to detect cell death (timeline as per Fig 2). **A-D**. Exemplar images of labelling (arrows). **E**. Decreasing mGluR5 signalling after TBI with 2-Methyl-6-(phenylethynyl)pyridine (MPEP) significantly increased neuronal cell death versus non-drug treated TBI larvae (p<0.0001). Increasing mGluR5 signalling after TBI with 2-Chloro-5-hydroxyphenylglycine (CHPG) significantly decreased CNS cell death versus non-drug treated TBI larvae (p<0.01). n = number of larvae, each experimental group was replicated twice. Bars represent the mean ± standard error. Data was analyzed using an ordinary one-way ANOVA with Tukey’s multiple comparison of means. Symbols indicate statistical significance: ^**^<0.01, ^***^<0.001, ^****^<0.0001, and ####<0.0001 versus all groups. Exact p-values and statistical details are listed in Supplemental Tables.

### Increased mGluR5 signalling in larvae with PTZ induced seizures results in decreased Tau aggregation

Increased neural activity, excitotoxicity, and seizures have all been shown to increase tauopathy in vivo^16–18,20–24^. Classically, increased mGluR5 signalling increases seizures but we and others have shown that, in certain disease contexts, agonizing mGluR5 can be seizure protective^37,46,47^. We predicted that mGluR5 signalling may be attenuating seizures after TBI as part of its neuroprotective mechanism. As a starting place based on our previous finding that seizures increase tauopathy after TBI^21^, we decided to induce seizures in our Tau biosensor larvae to assess if preventing seizure pathology is a part of mGluR5 mediated neuroprotection after TBI. Reducing mGluR5 signalling with MPEP in larvae experiencing seizures resulted in significantly higher Tau aggregation (Fig. 4, p<0.01), whereas increasing mGluR5 signalling with CHPG reduced Tau aggregation in seizing larvae (Fig. 4). Additionally, modulating mGluR5 signalling in uninjured larvae had no impact on Tau aggregation compared to no-treatment controls (Fig. 4), suggesting that pathophysiological changes are needed for mGluR5 activity to modulate tauopathy. These results suggest that the seizure activity caused by TBI may be the necessary (or one of many) pathophysiological changes needed for mGluR5 activity to mediate tauopathy.

**Figure 4.**
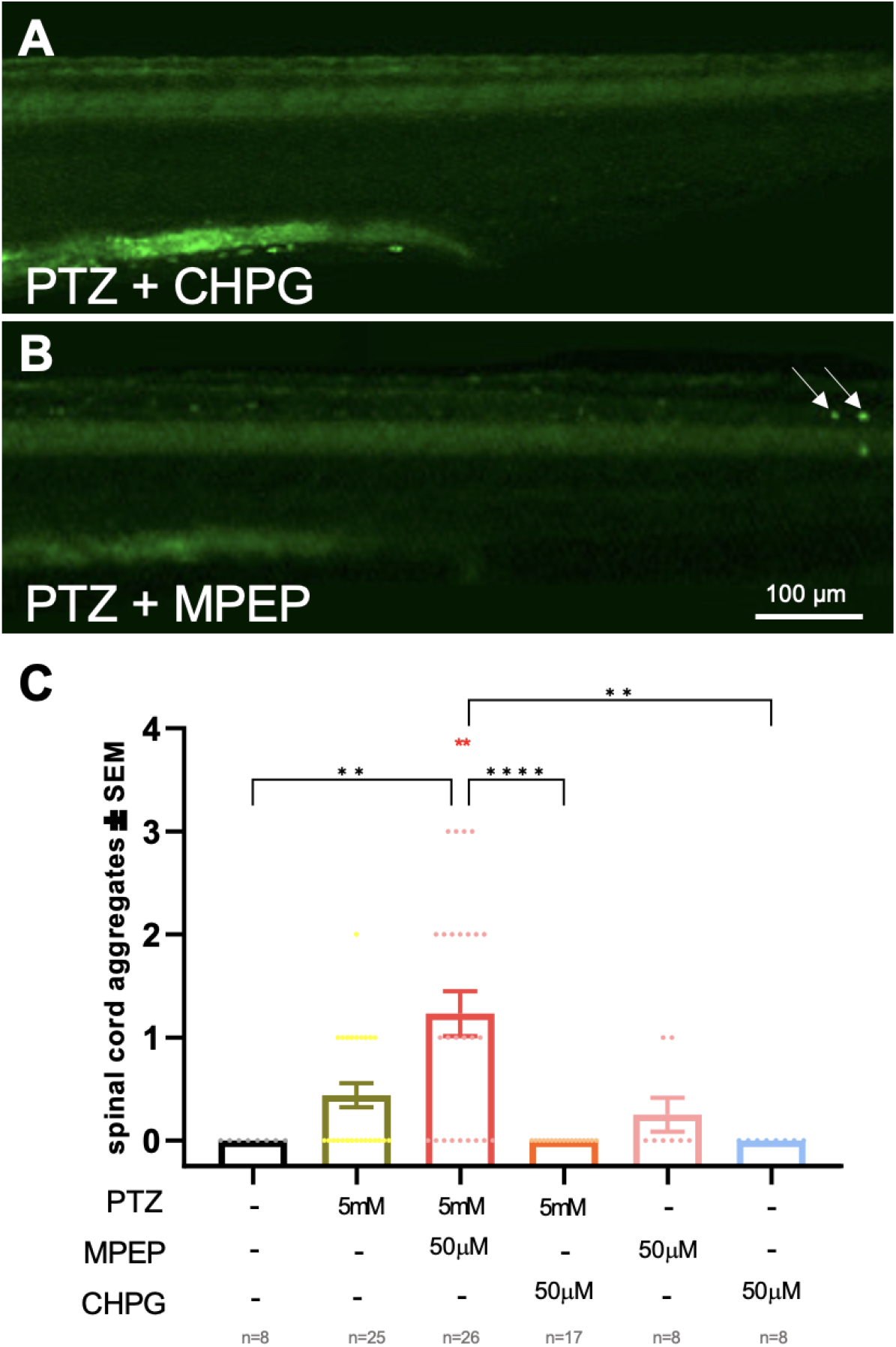
Increasing metabotropic glutamate receptor 5 (mGluR5) signaling in convulsing zebrafish larvae decreases Tau aggregation. At 3dpf larvae were treated with the convulsant pentylenetetrazole (PTZ) and/or 2-Methyl-6-(phenylethynyl)pyridine (MPEP) and/or 2-Chloro-5-hydroxyphenylglycine (CHPG) for 38 hours. At 7dpf the larvae were quantified via fluorescent microscopy by blinded observers. **A**,**B**. Exemplar images of tau aggregated induced by PTZ. **C**. PTZ caused a non-significant increase in GFP+ tau puncta compared to no treatment larvae. Decreasing mGluR5 signalling via MPEP in combination with PTZ further increased GFP+ tau puncta compared to no drug treated larvae (p<0.0001) and PTZ-only treated larvae (p<0.01). Decreasing mGluR5 signalling in combination with PTZ via CHPG prevented the formation of GFP+ tau puncta. n = number of animals. Bars represent the mean ± standard error. Data was analyzed using a Kruskal-Wallis test with Dunn’s multiple comparison. Symbols indicate statistical significance: ^**^ <0.01 and ^****^<0.0001. Exact p-values and statistical details are listed in Supplemental Tables.

### Activation of mGluR5 reduces tauopathy through the PI3K/Akt pathway following TBI

Activation of mGluR5 has been shown to be neuroprotective after TBI through signalling pathways such as the PI3K/Akt pathway for example, which differ from the toxic G_q_ protein signalling observed in AD^42,44,48–50^. Additionally, activation of the PI3K/Akt pathway has been shown to inactivate GSK3β, a kinase responsible for Tau hyperphosphorylation during disease progression^51,52^. We predicted that increased mGluR5 signalling was reducing tauopathy after TBI via neuroprotective downstream PI3K/Akt signalling. To assess if the neuroprotective effects of mGluR5 signalling after TBI are mediated by the PI3K/Akt pathway, we co-applied the mGluR5 agonist CHPG and PI3K/Akt antagonist LY294002. Inhibiting the PI3K/Akt pathway while activating mGluR5 signalling resulted in a significant increase in GFP+ Tau aggregates versus only activating mGluR5 (Fig. 5A and C, p<0.01). Additionally, inhibiting the PI3K/Akt pathway without increasing mGluR5 signalling resulted in a significant increase in GFP+ Tau aggregates at a lower dose of LY294002 (Fig. 5, p<0.05). These results support that protective mGluR5 signalling is mediated through the PI3K/Akt pathway since the drugs are able to counteract each other’s effects, suggesting pharmacological specificity toward the intended mGluR5-PI3K/Akt signalling pathway. Overall, these findings show that activating mGluR5 after TBI prevents tau pathology through downstream PI3K/Akt signalling.

**Figure 5.**
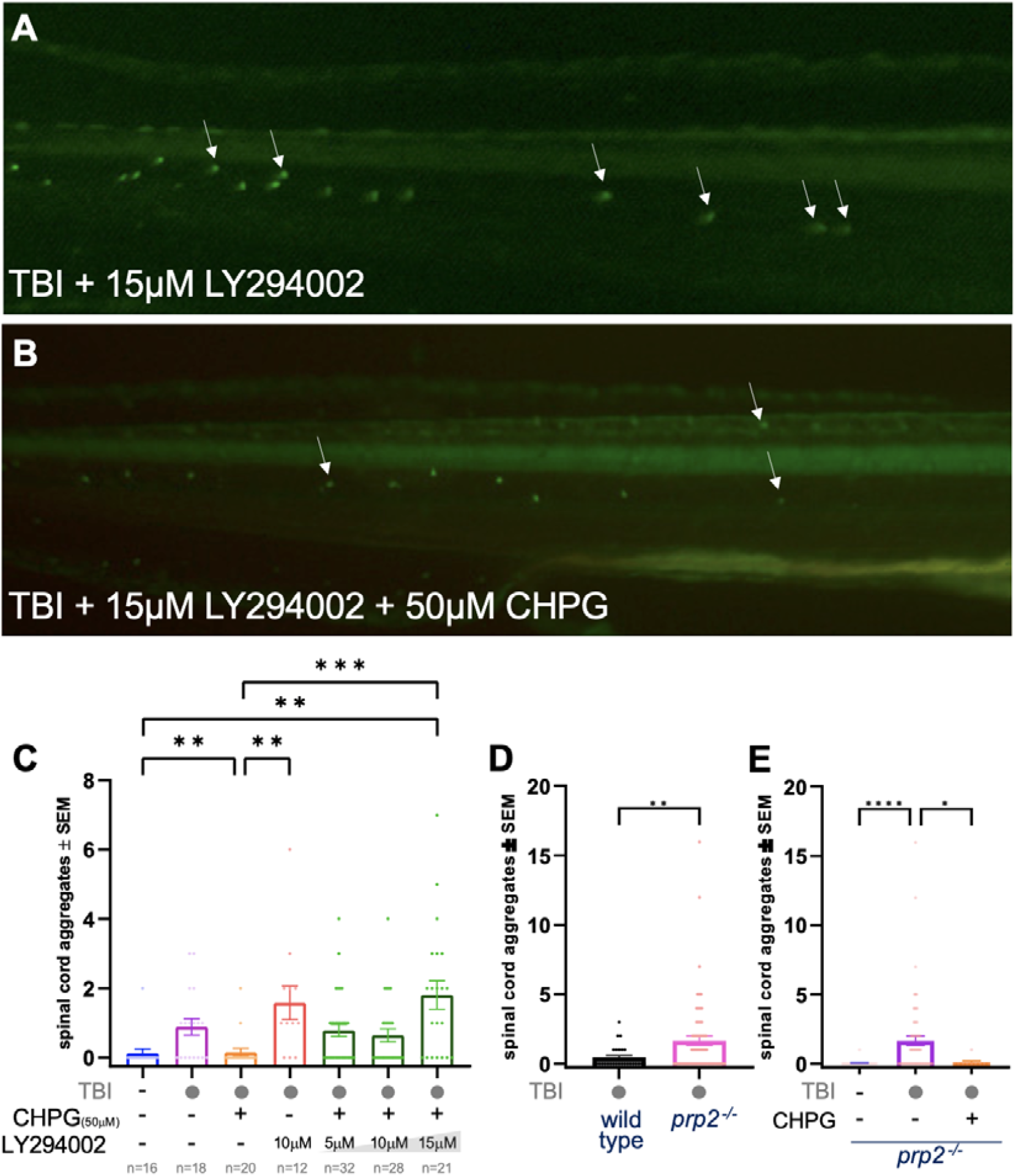
Activation of metabotropic glutamate receptor 5 (mGluR5) is protective through the PI3K/Akt pathway following TBI. Larvae were subjected to TBI at 3dpf and treated with drugs for 38 hours. At 7dpf the larvae were quantified via fluorescence microscopy by blinded observers, timeline as per Figure 2. **A**,**B**. Exemplar images of tau aggregated induced by TBI and LY294002. **C**. Inhibiting the PI3K/Akt pathway after TBI via LY294002 while mGluR5 signalling was activated with CHPG resulted in a removal of mGluR5 mediated tauopathy reduction as the dose of Ly294002 increased (p<0.01, 15µM LY294002 + 50µM CHPG + TBI vs 50uM CHPG + TBI). B/C. Representative images of larvae treated with 15µM LY294002 or 15µM LY294002 + 50µM CHPG after TBI, respectively. **D. Cellular Prion Protein (PrP**^C^**) is neuroprotective in zebrafish after TBI**. Wildtype and *prp2*^-/-^ larvae (lacking the homolog of mammalian *PRNP* that encodes PrP^C^) were subjected to TBI as per timeline in Figure 2. Each data point represents an individual animal. A. Loss of zebrafish *prp2* gene resulted in 3.7-fold increase in GFP+ Tau agrregates after TBI versus wild-type controls (p<0.01, Mann-Whitney test). **E**. *prp2*^−/−^ larvae did not display GFP+ Tau puncta without TBI. Additionally, increasing mGluR5 signalling with CHPG after TBI significantly reduced GFP+ Tau puncta (p<0.05, Kruskal-Wallis test with Dunn’s multiple comparison). Each data point represents an individual larva. n = number of animals, each experimental group was replicated at least twice. Bars represent the mean ± standard error. Data was analyzed using a Kruskal-Wallis test with Dunn’s multiple comparison. Symbols indicate statistical significance: ^*^<0.05 and ^**^<0.01. Exact p-values and statistical details are listed in Supplemental Tables.

### Loss of *prp2* increases GFP+ Tau aggregates in transgenic zebrafish after TBI

To further examine the opposing impacts of mGluR5 signalling in response to TBI versus Aβ oligomer injection, we next examined how loss of *prp2* impacts Tau pathology after TBI. Given that cellular prion protein knockout animals have been previously shown to be susceptible to brain injury^53,54^ and seizures^36,37,55^,we speculated that *prp2*^−/−^ zebrafish larvae would be more susceptible to tauopathy after TBI. To assess how loss of *prp2* effects tauopathy after TBI, we subjected *prp2*^−/−^ larvae^36^ to TBI while also altering mGluR5 signalling. Compared to wild-type larvae, *prp2*^−/−^ larvae had a 3.7-fold increase in GFP+ Tau puncta after TBI (Fig. 5D, p<0.01). Increasing mGluR5 signalling with CHPG after TBI reduced GFP+ Tau back to basal levels in *prp2*^−/−^ larvae (Fig. 5E, p<0.05). Decreasing mGluR5 signalling with MPEP after TBI in *prp2*^−/−^ larvae resulted in 100% mortality using doses that increased tauopathy in wild-type larvae. These results suggest that not only does loss of *prp2* lead to increased tauopathy after TBI, but it may also function to modulate downstream mGluR5 signalling levels given the hypersensitivity to decreased mGluR5 signalling.

### Increased metabotropic glutamate receptor 5 (mGluR5) signalling attenuates post traumatic seizure activity in *prp2*^-/-^ larvae

After establishing that loss of *prp2* and decreased mGluR5 signalling increases tauopathy after TBI, we next sought to verify how alterations to the PrP^C^-mGluR5 pathway impact post-traumatic seizures and test our hypothesis that the PrP^c^-mGluR5 pathway is a mechanistic link between post-traumatic seizures and tauopathy. To do so, we first measured if *prp2*^−/−^ zebrafish also exhibit seizure susceptibility after TBI, which could be a potential mechanism driving the increased tauopathy seen in *prp2*^−/−^ larvae after TBI. PrP^C^ knock-out animals, including *prp2*^−/−^ zebrafish, are well documented to have increased susceptibility to seizures when treated with convulsants^36,37,55^. To detect seizures, we used a previously validated behavioral assay^21,36,37,56^ to measure the seizure-like hyper-locomotory activity of 6dpf wild-type zebrafish larvae and *prp2*^−/−^ zebrafish larvae following TBI (Fig. 6). Post TBI, *prp2*^−/−^ larvae displayed a significant increase in seizure-like locomotor activity compared to control *prp2*^−/−^ larvae (p<0.01), control wild-type larvae (p<0.001) and post TBI wild-type larvae (p<0.0001) (Fig. 6A and B). Approximately 44% of post TBI *prp2*^−/−^ larvae displayed seizure-like activity reminiscent of stage I or II seizures, where larvae exhibit hyper-motility^57^.

**Figure 6.**
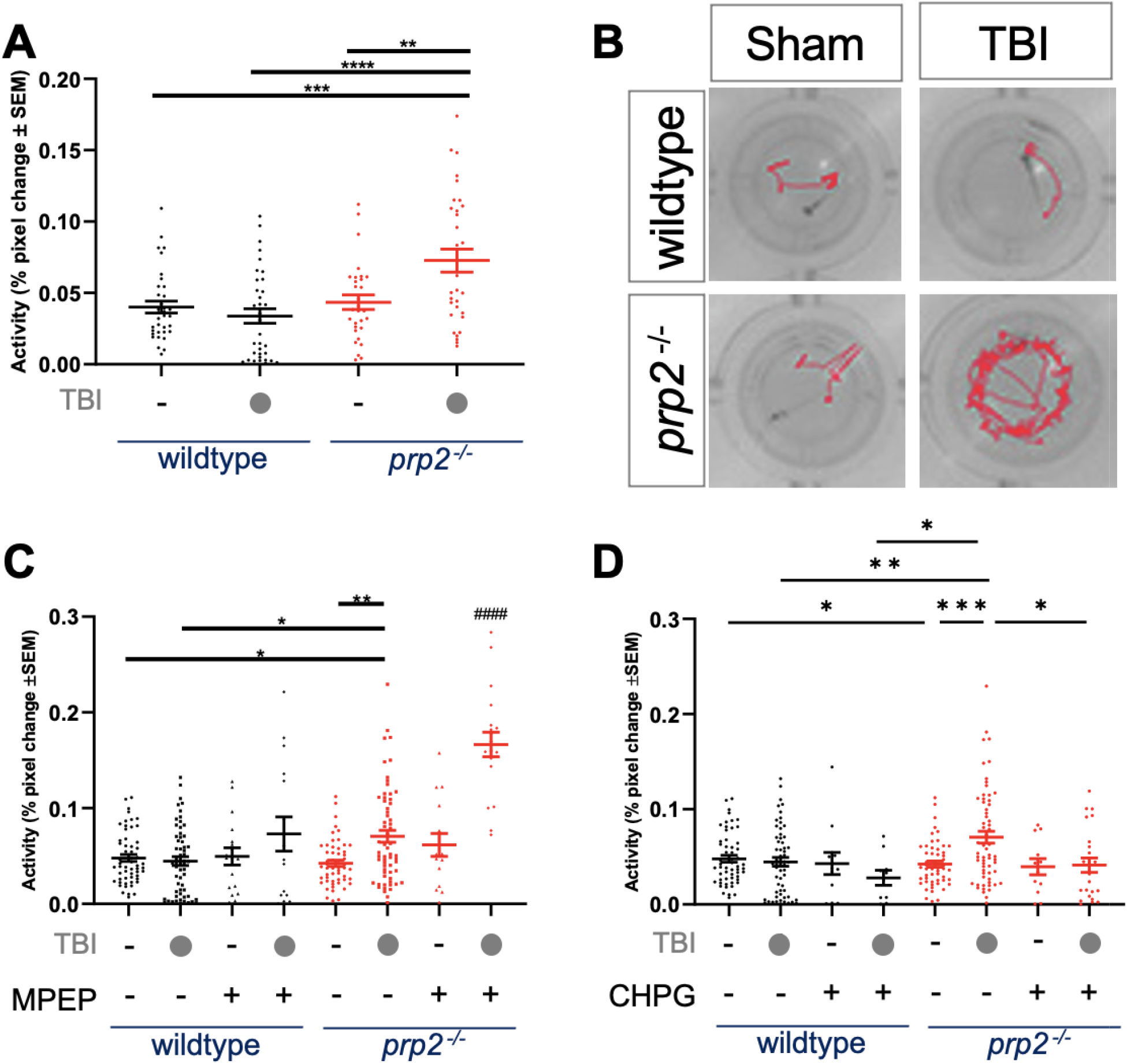
Increased metabotropic glutamate receptor 5 (mGluR5) signalling attenuates post-traumatic seizure-like behaviour in larvae lacking cellular prion protein (PrP^C^). Behavioral analysis of post-traumatic seizure activity in 6dpf wild-type and *prp2*^−/−^ larvae. Each dot is an individual animal, and horizontal bars indicate mean activity of the group +/-standard error of the mean (SEM). No TBI and TBI control data across panels A, C & D are the same data, groups were presented across multiple graphs for readability. **A**. *prp2* ^-/-^ larvae (red) with disrutped PrP^C^ displayed a significant increase in locomotor activity following traumatic brain injury (n = 24). **B**. Representative activity traces of larvae over a one-minute time frame, each imaging a single well of a 96-well plate. Seizure-like hyperactivity is induced by TBI in mutant larvae with disrupted PrP^C^ (bottom right) **C**. Inhibition of mGluR5 with 2-Methyl-6-(phenylethynyl)pyridine (MPEP) resulted in a significant increase in locomotor activity only in *prp2*^−/−^ larvae, only after inducing TBI (Red). When mGluR5 was inhibited in wild-type larvae no significant change in locomotor activity occurred with or without TBI (black) (n = 56, 59, 18, 16, 54, 62, 15, 19). **D**. Locomotor activity decreased to basal levels when mGluR5 was activated in *prp2*^−/−^ larvae via 2-Chloro-5-hydroxyphenylglycine (CHPG), whereas wild-type were not statistically affected (n = 56, 59, 12, 10, 54, 62, 12, 24). n = Number of animals, each experiment was replicated at least twice. Bars represent the mean ± standard error. Data was analyzed using Ordinary One-Way ANOVA and Tukey’s multiple comparison of means. Symbols indicate statistical significance: ^*^<0.05, ^**^<0.01, ^***^<0.001, ^****^<0.0001, and ####<0.0001 versus all groups. Exact p-values and statistical details are listed in Supplemental Tables.

To test if alterations to mGluR5 signalling impact post traumatic seizures, we next quantified post traumatic seizures while applying the mGluR5 antagonist MPEP. Wild-type larvae did not have any significant changes in locomotor activity after decreasing mGluR5 signalling, whereas *prp2*^−/−^ larvae showed approximately twice the level of seizure-like activity when mGluR5 was inhibited (p<0.001 vs. all treatments) (Fig 6C). For both genotypes MPEP treatment without TBI had a negligible impact on locomotor activity (Fig 6C), suggesting that some secondary factor mediated by TBI is needed to increase mGluR5-depndant seizure activity in *prp2*^−/−^ larvae. Antagonizing mGluR5 and disrupting PrP^C^ strongly potentiate each other’s effect on TBI-induced seizure-like activity.

This gave rise to our hypothesis that prion protein mutants are sensitized to post-traumatic seizures via downstream mGluR5 signalling; this predicts that agonizing mGluR5 signalling should rescue the prion mutant hypersensitivity. Thus, we applied the mGluR5 agonist CHPG after TBI. Post TBI CHPG treated *prp2*^−/−^ larvae showed a dose dependent decrease in locomotor activity returning to basal activity levels after 5µM treatment (Fig. 6D). Wild-type larvae did not display any significant changes to locomotor activity after CHPG treatment (Fig. 6D). These results are consistent with our results from Figure 6C, where alterations in locomotor activity due to disrupting the PrP^c^-mGluR5 pathway are only seen after TBI occurred.

Overall, these data suggest that inhibiting mGluR5 activity in *prp2*^−/−^ larvae exacerbates post traumatic seizures, whereas increasing mGluR5 activity leads to decreased post traumatic seizures. Moreover, the specific attribution of PTS susceptibility to a mechanism that involves PrP^C^ (Fig. 6A) is strongly supported by rescue of this phenotype via modulating mGluR5 (Fig 6D), a known PrP^C^ downstream effector.

## Discussion

Traumatic brain injuries (TBI) are a major cause of death and disability across all demographics having large scale impacts on individuals, families, and health systems^1–4^. While the direct injuries and consequences of TBI are important, relatively little is known about the pathological connection between TBI and subsequent tauopathies such as Alzheimer’s disease (AD) and chronic traumatic encephalopathy (CTE). Understanding the mechanisms in which TBI leads to tauopathy is key to developing treatment options and diagnostic tools for patients suffering from these tragic neurodegenerative diseases. In this study, we have built on our previous findings that post traumatic seizures lead to worse injuries and increased tauopathy^21^ with the goal of unravelling how post traumatic seizures lead to increased tauopathy. To do so, we used a combination of genetic and pharmaceutical methods to target the cellular prion protein (PrP^C^) and metabotropic glutamate receptor 5 (mGluR5) following TBI. This pathway was targeted for two primary reasons: 1) PrP^C^ knockout models display seizure susceptibility and increased seizure activity^35–37^; 2) mGluR5 and is downstream components have been implicated with Tau hyperphosphorylation, Tau aggregation and other pathologies in models of tauopathy^38,39,58–60^. Therefore, we hypothesized that post traumatic seizures are a mechanistic link to tau aggregation that acts in part through the PrP^C^-mGluR5 pathway.

## The PrP^C^-mGluR5 pathway is highly conserved in zebrafish larvae

It is well established that the PrP^C^-mGluR5 pathway is a conduit of toxic Aβ oligomer signalling during AD, but the conservation of Aβ mediated PrP^C^-mGluR5 pathology is relatively undefined in zebrafish specifically. Prior to this study, Özcan and colleagues showed that Aβ oligomers can bind and signal through the PrP^C^-mGluR5 pathway to mediate sleep-related behavioral changes in larval zebrafish. Using our transgenic Tau-biosensor zebrafish larvae, we further expanded the characterization of zebrafish PrP^C^-mGluR5 pathway conservation by measuring downstream tauopathy mediated by Aβ_1-42_ oligomers. Here, we showed that Aβ_1-42_ oligomers induce Tau pathology through PrP^C^-mGluR5 signalling through genetic and pharmacological manipulation. Aβ_1-42_ oligomer mediated pathology was reduced via pharmacological reduction of mGluR5 signalling displaying that the pathology is mediated through mGluR5 signalling (Figs. 1 and S1). Removal of zebrafish *prp2* prevented Aβ_1-42_ oligomer mediated pathology, confirming that zebrafish PrP2 is necessary for Aβ_1-42_ oligomer mediated mGluR5 pathology (Fig. 1B). Furthermore, pharmacological prevention of the PrP-mGluR5 interaction also reduced Aβ_1-42_ oligomer mediated tauopathy, meaning that the PrP-mGluR5 complex is needed for Aβ_1-42_ oligomer mediated tauopathy in our larval zebrafish model (Figs. 1 and S1). These findings validate the relevance of the larval zebrafish model for studying how/if the PrP^C^-mGluR5 pathway links TBI, seizures, and tauopathy.

## mGluR5 signalling is neuroprotective through the PI3K/Akt pathway after TBI

We next inspected the role of the PrP^C^-mGluR5 pathway on tauopathy after TBI. The potential implications of mGluR5 in pathological Tau formation is twofold. First, mGluR5 has been reported to mediate tau hyperphosphorylation in AD and other tauopathy models through downstream signaling of kinases and phosphatases^38,39,58–60^. Second, our recent discovery that increased post traumatic seizures exacerbate tau aggregation^21^ has led us to believe the functions of mGluR5 and interactions with PrP^C^ related to seizures make it a potential pathway linking post traumatic seizures and tauopathy. Our results showed that decreasing mGluR5 signalling via MPEP post-TBI increased Tau aggregation and neuronal cell death (Figs 2 and 3). Furthermore, these impacts were reciprocal when mGluR5 signalling was increased via CHPG (Figs 2 and 3) and these drugs can counteract each other’s effects (Figs 2 and 3) suggesting mGluR5 is specifically mediating this process. These results are opposing to what has previously been reported in AD models and our above findings of Aβ_1-42_ oligomer mediated tauopathy, where inhibiting mGluR5 and its downstream components prevents AD pathology^41,61–66^.

Although these results suggest the role of mGluR5 signalling after TBI is opposite to its role in Aβ oligomer mediated tauopathy, the results are consistent with previous TBI literature. Increased mGluR5 signalling mediated by the agonist CHPG reduces neuronal damage, apoptosis, edema, pro-inflammatory activation of microglia, and NMDA-mediated currents leading to neuroprotection and functional recovery in multiple neurotrauma models^42,43,45,48^. In these studies, mGluR5 signalling was shown to be neuroprotective through down streak ERK and/or PI3K/Akt signalling which activates these protective effects, possibly mediated through immune cell function^42–45,48^. In comparison, Aβ_1-42_ oligomers mediate pathology through toxic PrP^C^-mGluR5 G-protein signalling^38,39,62^. Thus, we considered that mGluR5 signalling is neuroprotective in our TBI model through alternate signalling methods compared to the Aβ_1-42_ oligomer mediated tauopathy we reported. We confirmed that mGluR5 signalling is neuroprotective through PI3K/Akt signaling by blocking the protective effects of mGluR5 mediated neuroprotection with the PI3K/Akt inhibitor LY2940002 (Fig. 5). Our conclusion that the neuroprotective effects of mGluR5 signalling are through alternate mechanisms to toxic Aβ_1-42_ oligomer-mediated mGluR5 signalling is further supported by our results that CHPG did not exacerbate tauopathy after Aβ_1-42_ oligomer injection (Fig. 1). If CHPG was enhancing toxic Aβ_1-42_ oligomer mediated PrP^C^-mGluR5 G-protein signalling then we would expect tauopathy to worsen.

The mechanism of reduced tauopathy after increased mGluR5 mediated PI3K/Akt signalling is interesting to consider and likely involves multiple factors. We found that decreasing mGluR5 signalling through MPEP did not create tauopathy without TBI (Fig. 2E), suggesting that pathological events are needed for mGluR5 signalling to influence tauopathy. We hypothesized that post-traumatic seizures are a mechanism that is regulated by PrP^C^-mGluR5 and drives tauopathy following TBI. To test this, we induced seizures in our larval zebrafish with a convulsant. Seizing zebrafish developed a non-significant amount of tauopathy but a concerted decrease in mGluR5 signalling significantly exacerbated tauopathy; Conversely, increasing mGluR5 signalling removed detectable tauopathy (Fig. 4). These results suggest that seizures are sufficient for mGluR5-mediated effects on tauopathy, meaning that increased mGluR5 signalling may be suppressing tauopathy driven by post-traumatic seizures. Although classic mGluR5 G-protein signalling causes increased neural activity, mGluR5 signalling has been shown to decrease seizures in a murine viral temporal lobe epilepsy model and Fragile-X syndrome model^46,47^. Additionally, mGluR5 signalling reduces glutamatergic activity and excitotoxicity in vitro by upregulating GluA2 subunits, specifically through PI3K/Akt signalling^67^.

Alternative mGluR5 functions that also need to be considered to understand the mechanism of neuroprotection against tauopathy include reducing Tau phosphorylation through glycogen synthase kinase 3 beta (GSKβ) inhibition and alterations in microglia function. Multiple studies have reported that PI3K/Akt signalling inhibits GSKβ, a major Tau phosphorylase^51,52,68^. In these studies, activation of PI3K/Akt via multiple mechanisms resulted in decreased Tau phosphorylation mediated by GSKβ suppression^51,68^ and increased cognitive function in a mouse model of AD^52^. Additionally, activating mGluR5 has been shown to attenuate microglial activity, which is neuroprotective and delays neurodegeneration after TBI^44,45,48,49^. The activation of mGluR5 is thought to be neuroprotective by reducing the inflammatory activation of microglia, instead causing microglia to adopt an anti-inflammatory phenotype^44^. Additionally, microglia have been shown to propagate tauopathy when activated in an inflammatory state^69–72^. Thus, mGluR5 mediated changes in microglia function through PI3K/Akt may also reduce tauopathy. These results and ours highlight the need to further establish the mechanisms of how the PrP^C^-mGluR5 pathway impacts tauopathy, especially when considering the design of targeted therapeutics since the pathway appears to have opposing functions on two closely related tauopathies in AD and CTE.

## PrP^C^ regulates seizures and mGluR5 function after TBI

To further query the role of PrP^C^-mGluR5 signalling on post-traumatic seizures and tauopathy we conducted TBI on *prp2*^−/−^ zebrafish. The loss of PrP^C^ has been shown to cause seizure susceptibility across multiple animal models^35–37^, in addition to susceptibility to brain damage following injuries^53,54,73,74^. Hence, we hypothesized that disrupting the PrP^C^-mGluR5 pathway through loss of *prp2* would increase tauopathy through increased post-traumatic seizures. Indeed, our data measuring GFP+ Tau aggregates showed that *prp2*^−/−^ larvae had a significant increase in detectable tauopathy after TBI (Fig. 5D). When mGluR5 signalling was increased *prp2*^−/−^ larvae had a significant reduction in tauopathy after TBI similar to wild-type larvae (Fig. 5E), indicating that the protective functions of mGluR5 exist without upstream PrP^C^. Interestingly, when mGluR5 signalling was decreased using doses effective in wild-type larvae there was complete mortality in the *prp2*^−/−^ larvae. These results suggest that zebrafish larvae lacking *prp2* are hypersensitive to decreased mGluR5 signalling after TBI. For example, upstream PrP^C^ may function as a regulator of downstream mGluR5 signalling meaning that removal of PrP^C^ decreases endogenous signalling or mGluR5’s sensitivity to ligands. PrP^C^ has been shown to regulate PI3K/Akt signalling levels, where loss of PrP^C^ decreases signalling levels and overexpression of PrP^C^ increases signalling levels^54,75,76^. Even so, it is unclear whether PI3K/Akt signalling regulated by PrP^C^ is through downstream mGluR5 activity, so further studies are needed.

We next examined the contributions of PrP^C^-mGluR5 activity on post-traumatic seizures as a possible mechanism responsible for the increased tauopathy seen with loss of zebrafish PrP2. Our results showed that *prp2*^−/−^ larvae do indeed display a susceptibility to post traumatic seizures after TBI (Fig. 6A). We further examined PrP^C^ susceptibility to post traumatic seizures by targeting its downstream counterpart mGluR5 via pharmaceuticals. Our results showed that inhibiting mGluR5 activity in *prp2*^−/−^ larvae significantly exacerbated post traumatic seizures, and conversely post traumatic seizures can be mitigated by activating mGluR5 (Fig. 6C). These results provide further evidence that PrP^C^ may be acting as a regulator of downstream mGluR5 activity after TBI since *prp2*^−/−^ were hypersensitive to the effects of mGluR5 signalling on post-traumatic seizures. Based on our findings, PrP^C^ appears to dampen seizures after TBI possibly through the regulation of downstream mGluR5 activity. Moreover, when the PrP^C^-mGluR5 pathway is disturbed via disruption of PrP^C^ tauopathy is increased after TBI. These results support our hypothesis that the PrP^C^-mGluR5 pathway is a mechanism whereby TBI leads to seizures and subsequent tauopathy. This is further supported by our data where decreasing mGluR5 signalling in seizing larvae exacerbates tauopathy (Fig. 4), which suggests that seizure pathology in combination with mGluR5 disruption leads to tauopathy.

Overall, these results merit consideration when both PrP^C^ and mGluR5 are potential targets for AD therapies. The contributions of the PrP^C^-mGluR5 pathway to AD pathology are well defined and therapies targetting this pathway are promising, but such a therapeutic approach may have negative consequences for patients with a history of TBI. Our findings align with previous results outlining the neuroprotective functions of PrP^C^ and mGluR5 activity after TBI while further detailing how these functions impact TBI mediated tauopathy. The PrP^C^-mGluR5 pathway should be considered for further research as a therapeutic target for post traumatic seizures and tauopathy after TBI.

## Limitations

Although our larval zebrafish TBI method has many practical and ethical advantages, the limitations need to be taken into consideration when expanding our results to human physiology. Firstly, zebrafish have a duplicate genome so the complexity of having two versions of the proteins/genes studied in this manuscript needs to be considered with the functions and interactions of these proteins. Our results display that our *prp2*^−/−^ mutants have similar phenotypes to other PrP^C^ knockout models and our results from modulating mGluR5 activity with pharmaceuticals are consistent with previous literature, but it is still possible that the paralogs of mGluR5 and PrP^C^ genes have unique functions and interactions which do not completely reflect human biology. Secondly, the logistics of our zebrafish TBI model has limitations since we use larval zebrafish over a relatively short amount of time. Although larval zebrafish have a functioning central nervous system, they are still undergoing development which means that their unique biology might be most reflective of a human fetus. Additionally, our studies were conducted over the time frame of a week whereas the development of neurodegenerative tauopathies tends to take place over years so even though our system shows acute changes in tauopathy it does not explore factors that influence the long-term development of disease. Yet, our approach may have good fidelity in representing events immediately after TBI, at the launch of pathophysiology, and this is an attractive/practical therapeutic window in neurotrauma patients. Moreover, our Tau4R-GFP metric of tau aggregation is not mainstream for the tauopathy field, such that further work assessing tau biochemistry and phosphorylation is warranted; though we note this reporter has been thoroughly validated, responds as predicted to external insults, and responds appropriately across broad doses of intervention (doses of injury, doses of compounds)^21,78–80^.

## Conclusion

The pathological mechanisms and causes of tauopathies remain mysterious, and no viable treatments exist for these tragic neurodegenerative diseases. Here, we have found that PrP^C^ and mGluR5 signalling is protective against the formation of Tau aggregates after TBI. Increasing the activity of mGluR5 after TBI decreases tauopathy through the PI3K/Akt pathway.

Additionally, *prp2*^−/−^ zebrafish are susceptible to post traumatic seizures and tauopathy after TBI, possibly through the loss of PrP^C^ ‘s function as a regulator of downstream mGluR5 activity. TBI appears to be needed for mGluR5 to impact tau aggregation suggesting that a physiological disruption caused by TBI is needed before mGluR5 is involved in tauopathy. Our initial results suggest that seizures may be (one of) the factor needed for decreased mGluR5 activity to contribute to tau aggregation. These data provide a step towards understanding mechanisms of how post traumatic seizures contribute to TBI pathomechanisms, at least in an animal model. These results need to be expanded to understand how the other components of the PrP^C^-mGluR5 pathway are involved in these events and how suppressing mGluR5 activity leads to increased Tau aggregation, and how these manipulations would impact longer-term disease progression in a mammalian context.

## Methods

### Animal ethics and Zebrafish Husbandry

Zebrafish were bred and maintained following protocol AUP00000077 approved by the Animal Care and Use Committee: Biosciences at the University of Alberta, operating under the guidelines of the Canadian Council of Animal Care. The fish were raised and maintained within the University of Alberta fish facility under a 14/10 light/dark cycle at about 28°C as previously described^77^. Adult animals were bred to generate larval fish. All experiments used larval fish, and thus number of animals used in not applicable. Sex of larval zebrafish is unknowable. Animals were euthanized for regular colony maintenance by bath exposure to excess MS-222 (Sigma E10521). Adult animals were monitored twice daily with Veterinarian oversight, and their environmental conditions were monitored continuously by remote sensors.

Zebrafish used were AB strain, and included *prp2* mutants with a loss-of-function mutation in the homolog of PrP^C^ (ZDB-GENE-041221-3, *prnprs3*)^55^. Transgenics carrying the tauopathy reporter Tau4R-GFP, which is a fusion of GFP to the human Tau repeat region [*Tg(eno2:Hsa*.*MAPT_Q244-E372*−*EGFP)ua3171* larvae (ZFIN ID: ZDB-ALT-211005-6; research resource identifier (RRID) number RRID:ZFIN_ZDB-ALT-211005-6)], are co-expressed with full-length human tau (no GFP physically linked; *Tg(eno2:hsa*.*MAPT-ires-egfp)Pt406* larvae (ZFIN ID: ZDB-ALT-080122–6; RRID:ZFIN_ZDB-ALT-080122-6).

Animals were excluded if they exhibited overt morphological phenotypes. Larvae were randomly plated in dishes for subsequent treatment and analysis. Observers were blinded after treatments were applied and prior to observing the animals or quantifying outcomes. The study design used previously-validated outcome measures (tau aggregation, cell death and seizure-like behaviour) as endpoints.

### Amyloid beta 1-42 oligomer formation

Amyloid beta Aβ_1-42_ monomers and oligomers were prepared following previously published protocols^40^. Briefly, Aβ_1-42_ peptides (rPeptide, #A-1163-1) were dissolved in DMSO (100µM concentration) and kept at room temperature for 12 minutes. The peptide solution was then sonicated and stored. To prepare the Aβ solution for injections, 1 µL of the stock solution was diluted in PBS and the diluted solution was then incubated at 37°C for 24 hours to form oligomers. In experiments applying Aβ monomers, the incubation step was skipped and freshly thawed Aβ solution was diluted and used immediately in the subsequent steps. Finally, a 1:10 serial dilution using PBS was used to obtain a final concentration of 10nM Aβ.

### *Amyloid beta* Aβ_1-42_ *oligomer injection*

Aβ oligomers were injected into the hindbrain ventricle of 3dpf zebrafish larvae using previously described methods^21^. 3 days post fertilization zebrafish larvae were anesthetized with 4% tricaine and placed in small holes poked into a 1% agarose-coated petri dish, oriented so that their brain ventricle was accessible for injection. Injections were done under the view of a dissecting microscope, with pulled capillary tubes housed by a micromanipulator. The injection volume of 10nM Aβ solution was calculated and calibrated to be 1 nL. All injection had red dextran fluorescent dye mixed into the solution to confirm injection. Control injected groups received PBS mixed with red dextran fluorescent dye.

### Bath application of drugs for Tau aggregate quantification

Tau biosensor larvae were treated with drugs at 3dpf, 2-3 hours after Aβ injection or TBI and time-matched in sham controls. Larvae were left in the drug media for approximately 38-44 hours before the media was washed out, and tauopathy was quantified at 4 days post-injury. Doses for MPEP (2-Methyl-6-(phenylethynyl)pyridine (Tocrisis Bioscience, #1212) were chosen based on our previous publication^37^ and a dose response experiment (Fig. 2). For CHPG (2-Chloro-5-hydroxyphenylglycine, Hello Bio, #HB0034), a dose response experiment based on the effective doses of MPEP was used to determine appropriate concentrations (Fig. 2). Doses of LY294002 (MedChemExpress HY-10108) were chosen from a dose response experiment (Fig. 5). Lastly, for BMS-984923 (MedChemExpress HY-122559) a dose response experiment was conducted to measure the efficacy and toxicity of the drug in larval zebrafish (Supplemental Fig. S1B). At 7dpf Tau biosensor larvae were visualized using the methods previously described to quantify GFP+ Tau aggregates^21^.

### Analysis of Tau GFP+ aggregates

The larvae that were injured at 3dpf were quantified at 7dpf using a Leica M165 dissecting microscope. GFP+ puncta were quantified by blinded observers using methods previously described^21^. All larvae were anesthetized at the time of quantification using tricaine.

### Induction of TBI on larval zebrafish

To induce TBI we used methods similar to those previously described^21,79,80^. In short, 12-18 larvae were placed in a 20mL syringe (Becton Dickinson #302830) with 1mL of water (E3 media) before being closed off with a stopper valve. Larvae were 3 dpf for Tau quantification and 6 dpf for seizure quantification. The syringe was placed vertically in a tube holder clamp at the bottom of a 48” tube. A 300g weight was dropped from the top of the tube, producing pressure waves with approximately 450 kPa maximal pressure. After each drop, the larvae were repositioned by opening the plunger and ensuring any larvae pushed into the syringe tip were moved back into the barrel. This procedure was repeated three times per experiment to minimize inter-individual variability in the injury event. After injury, larvae were returned to petri dishes with fresh media and used for further analysis.

### Immunohistochemistry

Whole-mount immunostaining of Activated-Caspsase-3 was performed on zebrafish larvae following previously described protocols (DuVal et al. 2014). 7dpf larvae were fixed in 4% paraformaldehyde overnight at 4C. Larvae were washed three times in 0.1M PO_4_ + 5% sucrose solution, followed by a 1% tween + H_2_O wash, and −20C acetone wash. A 1-hour blocking step was performed by washing larvae with PBS3+ containing 10% goat serum. The larvae were then incubated with primary antibody in 2% goat serum PBS3+ overnight, followed by 3 PBS3+ washes, and lastly incubated with secondary antibody in 2% goat serum PBS3+. The primary antibody used was polyclonal Anti-Active-Caspase-3 (BD Biosciences Cat# 559565, RRID:AB_397274) at 1:500 dilution. The secondary antibody used was Alexafluor 488 anti-rabbit at 1:250 dilution (Invitrogen).

Example images of immunostained larvae were taken using a (get model) confocal microscope. The larvae were mounted on glass slides using 2% high-melting point agarose.

### Behavioral assay for detection of seizure-like movements

The behavioral tracking software EthoVision® XT-11.5 (Noldus, Wageningen, Netherlands; RRID:SCR_000441) was used to quantify seizure-like phenotypes using methods from our recent publications^21,36,37,56^. Briefly, 6dpf wild-type and *prp2*^−/−^ larvae subjected to TBI 3x with a 300g weight were immediately placed into 96 well plates with 100µL of E3 embryo media, healthy larvae of both genotypes were included as controls. In experiments where drugs were included, the larvae were placed with 50µL of E3 embryo media, and 50µL of the drug was added at 2x working concentration approximately 20 minutes after TBI. The drugs used in this study were MPEP and CHPG, as described above. Behavioral activity recordings were started 45 minutes after the first TBI event for each experiment. Once placed, the larvae were recorded by an overhead camera under infrared light using EthoVision® XT-11.5, where 30 minutes of activity was collected for quantification as mean activity.

Seizure phenotypes are defined as hypermotility for stage I and II seizures, while stage III seizures are arrhythmic convulsions that appear as decreased movement in this assay since they display as epileptic convulsions^36,37^. Activity in EthoVision XT-11.5 is calculated as the % of pixel changes within each specific well between changes of frames during recording (recorded at 25fps). The values reported appear quite small given the tiny size of zebrafish larvae, but the assay is well tuned to detecting robust zebrafish movement in a reproducible and accurate manner.

### Statistics

All statistics were performed using GraphPad Prism 9 (GraphPad, San Diego CA; RRID:SCR_002798). Each experiment had at least two replicates and the sampling unit in each figure were individual larvae. Outliers were excluded from EthoVision® XT-11.5 data due to the rare mis-tracking of larval zebrafish resulting in locomotor results that are inaccurate and if values of zero were determined to be dead larvae. The experimenters were blinded in all experiments with manual quantifications of Tau aggregates, where outliers were not excluded.

Sample sizes were not determined *a priori* and instead were guided by previous works using these endpoints^21,78–80^. No data normalization was used. The data was not normally distributed, so analysis of experiments with more than two groups was completed using either Kruskal-Wallis test with Dunn’s multiple comparison or Ordinary One-Way ANOVA with Tukey’s multiple comparison of means post hoc test, as indicated. Analysis of two groups was completed using Mann-Whitney test. Exact p-values and statistical details are listed in Supplemental Tables.

## Abbreviations

Aβ: amyloid beta
AD: Alzheimer Disease
AKT: protein kinase B (name from AK thymoma retrovirus)
BMS: BMS-984923
CHPG: 2-Chloro-5-hydroxyphenylglycine
CTE: chronic traumatic encephalopathy
GFP+: green fluorescent protein-positive
GSK3β: glycogen synthase kinase 3 beta
mGluR5: metabotropic glutamate receptor 5
MPEP: 2-Methyl-6-(phenylethynyl)pyridine
PI3K: Phosphoinositide 3-kinase
*prp2*: zebrafish gene homolog of PrP^C^
PrP^C^: cellular prion protein
RRID: Research resource identifier
TBI: Traumatic Brain Injury
Tau-4R: four-repeat region of the human Tau protein

## Acknowledgments

Edward Burton provided their transgenic zebrafish line, a component of our dual transgenic zebrafish (Tau4R-GFP and human Tau 0N2R). We appreciate the care of University of Alberta Animal Care staff and Veterinarians. HA would like to thank the Deanship of Postgraduate Studies and Scientific Research at Majmaah University for supporting her work.

## Author contributions

LFL designed the project, acquired data, analyzed and interpreted data, and wrote the manuscript. JN, MJK and TG collected data and helped with data interpretation. MJK curated data, analyzed data, and assisted with visualization. HA contributed to conceptualization and edited the manuscript. WTA was responsible for conception and designing the project, interpreting and visualizing the data, supervision, funding, and writing the manuscript. All authors reviewed and edited the manuscript.

## Data Availability

Data will be made available upon request.

---

**Supplemental Figure S1.**
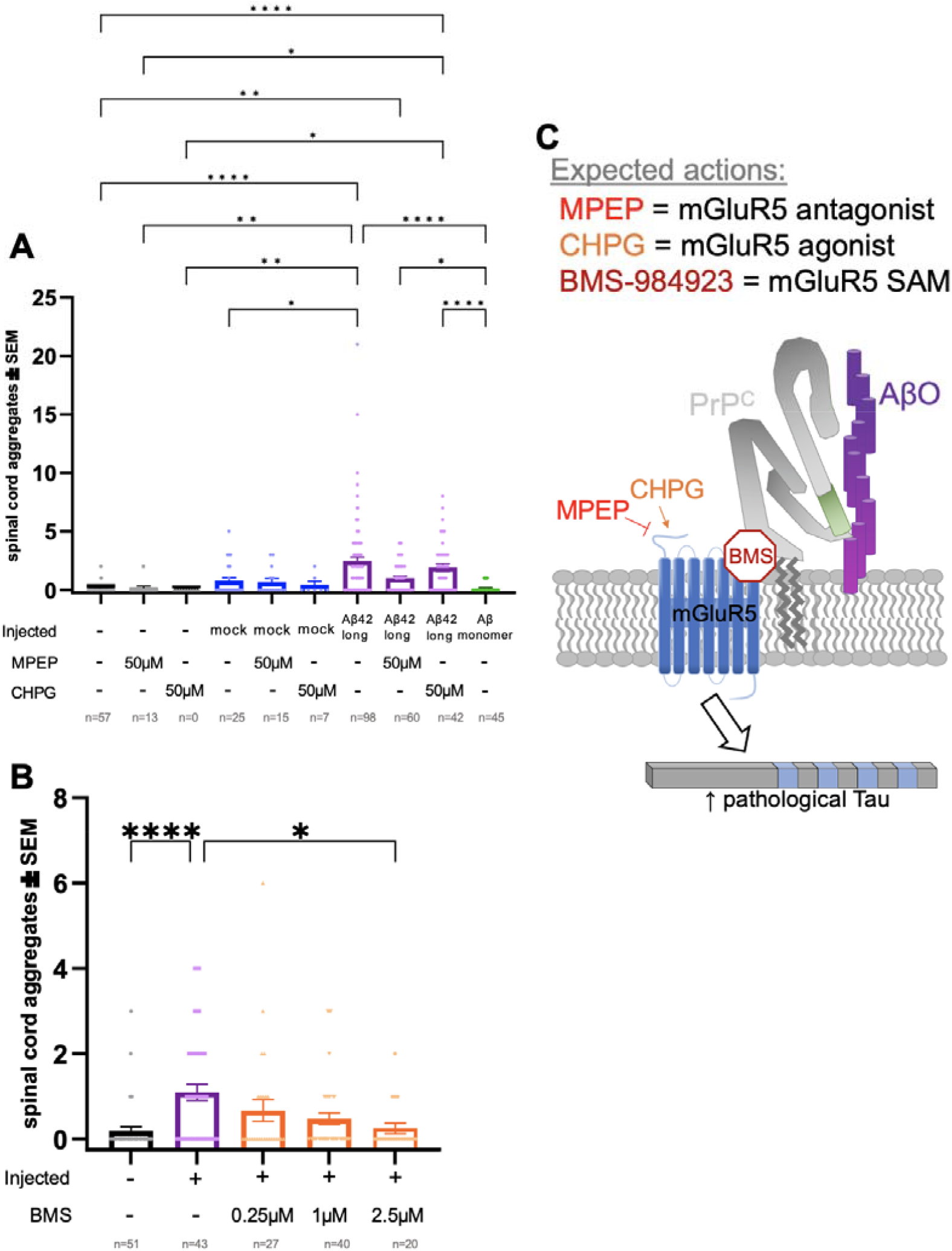
Injecting human amyloid beta_1-42_ oligomers *(AβO)* and manipulating mGluR5-PrP^C^ alters tau aggregation in a fashion highly reminiscent of mammalian systems. Several of these data sets are displayed in Figure 1 and repeated here for reference. **A**. Injection of AβO significantly increased GFP+ tau puncta vs un-injected (p<0.001) and mock injected (p<0.01) controls, whereas injecting monomers had no discernible impact. Decreasing mGluR5 signaling via MPEP decreased detectable tau puncta after AβO injection (p<0.001). Impacts of CHPG were negligible. **B**. BMS-984923 reduces the impact of AβO injection in a dose-dependent manner (p<0.05 injected control vs injected + 2.5µM BMS). **C**. Schematic summary of the expected actions of the compounds used. Broadly, AβO injection leads to tau aggregation via binding to cellular prion protein (PrP^C^) then signalling thru mGluR5 and downstream kinases to phosphorylate tau. 2-Methyl-6-(phenylethynyl)pyridine (MPEP) agonizes mGluR5. 2-Chloro-5-hydroxyphenylglycine (CHPG) is an mGluR5 antagonist. BMS-984923 is a silent allosteric modulator (SAM) that blocks mGluR5’s interaction with cellular PrP^C^-AβO complexes. n = the number of larvae in each group, each experimental group was replicated at least twice. Each dot is an individual animal. Bars represent the mean ± standard error. Data were analyzed using a Kruskal-Wallis test with Dunn’s multiple comparison. Symbols indicate statistical significance: ^*^ <0.05, ^**^<0.01, and ^****^<0.0001. Exact p-values and statistical details are listed in Supplemental Tables.

