## Supplemental Figure & Tables for "Seizures and tauopathy following neurotrauma are mediated by prion protein and metabotropic glutamate receptor 5"

Contents include:

- Supplemental Figure S1, accompanies Figure 1.
- Tables of statistical tests that accompany the manuscript's figures.

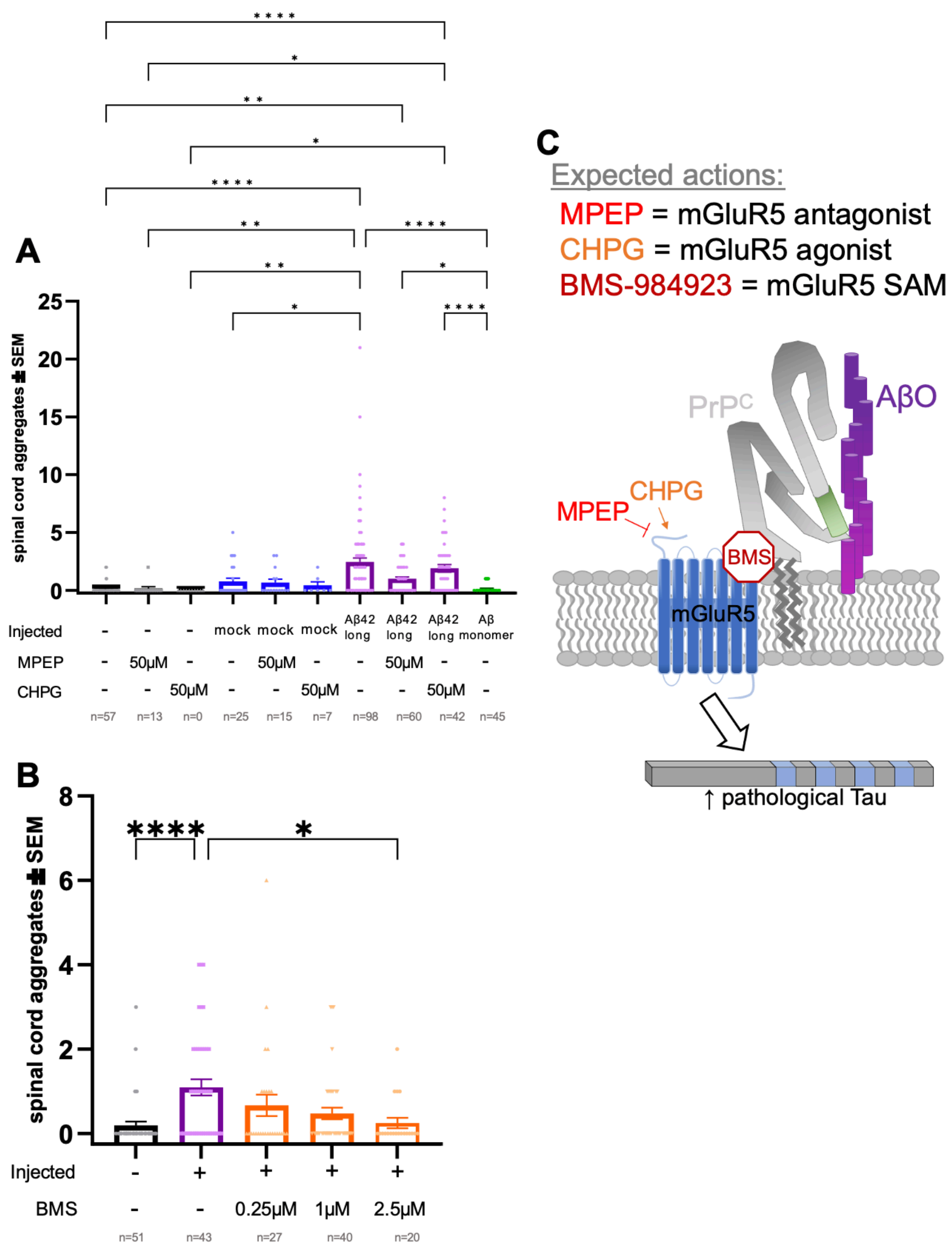

**Supplemental Figure S1. Injecting human amyloid beta$_{1\text{-}42}$ oligomers** (AβO) **and manipulating mGluR5-PrP$^{C}$ alters tau aggregation in a fashion highly reminiscent of mammalian systems.** Several of these data sets are displayed in Figure 1 and repeated here for reference. **A.** Injection of AβO significantly increased GFP+ tau puncta vs un-injected ($p<0.001$) and mock injected ($p<0.01$) controls, whereas injecting monomers had no discernible impact. Decreasing mGluR5 signaling via MPEP decreased detectable tau puncta after AβO injection ($p<0.001$). Impacts of CHPG were negligible. **B.** BMS-984923 reduces the impact of AβO injection in a dose-dependent manner ($p<0.05$ injected control vs injected + 2.5μM BMS). **C.** Schematic summary of the expected actions of the compounds used. Broadly, AβO injection leads to tau aggregation via binding to cellular prion protein ($PrP^C$) then signalling thru mGluR5 and downstream kinases to phosphorylate tau. 2-Methyl-6-(phenylethynyl)pyridine (MPEP) agonizes mGluR5. 2-Chloro-5-hydroxyphenylglycine (CHPG) is an mGluR5 antagonist. BMS-984923 is a silent allosteric modulator (SAM) that blocks mGluR5's interaction with cellular $PrP^C$-AβO complexes. n = the number of larvae in each group, each experimental group was replicated at least twice. Each dot is an individual animal. Bars represent the mean ± standard error. Data were analyzed using a Kruskal-Wallis test with Dunn's multiple comparison. Symbols indicate statistical significance: * <0.05, **<0.01, and ****<0.0001. Exact p-values and statistical details are listed in Supplemental Tables.

### Analysis associated with Figure 2E

| | Kruskal-Wallis test<br>ANOVA results | |
|---|---|---|
| 1 | Table Analyzed | SCA_TBI_MPE |
| 2 | | |
| 3 | **Kruskal-Wallis test** | |
| 4 | P value | <0.0001 |
| 5 | Exact or approximate P value? | Approximate |
| 6 | P value summary | **** |
| 7 | Do the medians vary signif. (P < 0.05)? | Yes |
| 8 | Number of groups | 9 |
| 9 | Kruskal-Wallis statistic | 74.35 |
| 10 | | |
| 11 | **Data summary** | |
| 12 | Number of treatments (columns) | 9 |
| 13 | Number of values (total) | 222 |

### Analysis associated with Figure 2E

| | **Kruskal-Wallis test**<br>Multiple comparisons | | | | | |
|---|---|---|---|---|---|---|
| 1 | Number of families | 1 | | | | |
| 2 | Number of comparisons per family | 36 | | | | |
| 3 | Alpha | 0.05 | | | | |
| 4 | | | | | | |
| 5 | **Dunn's multiple comparisons test** | **Mean rank diff.** | **Significant?** | **Summary** | **Adjusted P Value** | |
| 6 | No TBI vs. TBI | -52.72 | Yes | **** | <0.0001 | A-B |
| 7 | No TBI vs. TBI + DMSO | -66.88 | No | ns | 0.1768 | A-C |
| 8 | No TBI vs. TBI + 3mM MPEP | -30.45 | No | ns | >0.9999 | A-D |
| 9 | No TBI vs. TBI + 10mM MPEP | -68.75 | Yes | * | 0.0270 | A-E |
| 10 | No TBI vs. TBI + 30mM MPEP | -71.30 | Yes | **** | <0.0001 | A-F |
| 11 | No TBI vs. TBI + 50mM MPEP | -99.93 | Yes | **** | <0.0001 | A-G |
| 12 | No TBI vs. TBI + 100mM MPEP | -91.96 | Yes | **** | <0.0001 | A-H |
| 13 | No TBI vs. 50mM MPEP | -10.39 | No | ns | >0.9999 | A-I |
| 14 | TBI vs. TBI + DMSO | -14.15 | No | ns | >0.9999 | B-C |
| 15 | TBI vs. TBI + 3mM MPEP | 22.27 | No | ns | >0.9999 | B-D |
| 16 | TBI vs. TBI + 10mM MPEP | -16.03 | No | ns | >0.9999 | B-E |
| 17 | TBI vs. TBI + 30mM MPEP | -18.58 | No | ns | >0.9999 | B-F |
| 18 | TBI vs. TBI + 50mM MPEP | -47.21 | Yes | * | 0.0158 | B-G |
| 19 | TBI vs. TBI + 100mM MPEP | -39.24 | No | ns | 0.6365 | B-H |
| 20 | TBI vs. 50mM MPEP | 42.34 | No | ns | >0.9999 | B-I |
| 21 | TBI + DMSO vs. TBI + 3mM MPEP | 36.42 | No | ns | >0.9999 | C-D |
| 22 | TBI + DMSO vs. TBI + 10mM MPEP | -1.879 | No | ns | >0.9999 | C-E |
| 23 | TBI + DMSO vs. TBI + 30mM MPEP | -4.429 | No | ns | >0.9999 | C-F |
| 24 | TBI + DMSO vs. TBI + 50mM MPEP | -33.05 | No | ns | >0.9999 | C-G |
| 25 | TBI + DMSO vs. TBI + 100mM MPEP | -25.08 | No | ns | >0.9999 | C-H |
| 26 | TBI + DMSO vs. 50mM MPEP | 56.49 | No | ns | >0.9999 | C-I |
| 27 | TBI + 3mM MPEP vs. TBI + 10mM MPEP | -38.30 | No | ns | >0.9999 | D-E |
| 28 | TBI + 3mM MPEP vs. TBI + 30mM MPEP | -40.85 | No | ns | 0.9929 | D-F |
| 29 | TBI + 3mM MPEP vs. TBI + 50mM MPEP | -69.48 | Yes | ** | 0.0046 | D-G |
| 30 | TBI + 3mM MPEP vs. TBI + 100mM MPEP | -61.51 | No | ns | 0.0992 | D-H |
| 31 | TBI + 3mM MPEP vs. 50mM MPEP | 20.06 | No | ns | >0.9999 | D-I |

### Analysis associated with Figure 2E

| | **Kruskal-Wallis test** Multiple comparisons | | | | | | |
|---|---|---|---|---|---|---|---|
| 32 | TBI + 10mM MPEP vs. TBI + 30mM MPEP | -2.550 | No | ns | >0.9999 | E-F | |
| 33 | TBI + 10mM MPEP vs. TBI + 50mM MPEP | -31.18 | No | ns | >0.9999 | E-G | |
| 34 | TBI + 10mM MPEP vs. TBI + 100mM MPEP | -23.21 | No | ns | >0.9999 | E-H | |
| 35 | TBI + 10mM MPEP vs. 50mM MPEP | 58.37 | No | ns | >0.9999 | E-I | |
| 36 | TBI + 30mM MPEP vs. TBI + 50mM MPEP | -28.63 | No | ns | >0.9999 | F-G | |
| 37 | TBI + 30mM MPEP vs. TBI + 100mM MPEP | -20.66 | No | ns | >0.9999 | F-H | |
| 38 | TBI + 30mM MPEP vs. 50mM MPEP | 60.92 | No | ns | 0.8317 | F-I | |
| 39 | TBI + 50mM MPEP vs. TBI + 100mM MPEP | 7.969 | No | ns | >0.9999 | G-H | |
| 40 | TBI + 50mM MPEP vs. 50mM MPEP | 89.54 | Yes | * | 0.0266 | G-I | |
| 41 | TBI + 100mM MPEP vs. 50mM MPEP | 81.57 | No | ns | 0.1392 | H-I | |
| 42 | | | | | | | |
| 43 | **Test details** | **Mean rank 1** | **Mean rank 2** | **Mean rank di** | **n1** | **n2** | **Z** |
| 44 | No TBI vs. TBI | 61.70 | 114.4 | -52.72 | 51 | 62 | 4.728 |
| 45 | No TBI vs. TBI + DMSO | 61.70 | 128.6 | -66.88 | 51 | 7 | 2.813 |
| 46 | No TBI vs. TBI + 3mM MPEP | 61.70 | 92.15 | -30.45 | 51 | 17 | 1.843 |
| 47 | No TBI vs. TBI + 10mM MPEP | 61.70 | 130.5 | -68.75 | 51 | 10 | 3.370 |
| 48 | No TBI vs. TBI + 30mM MPEP | 61.70 | 133.0 | -71.30 | 51 | 25 | 4.951 |
| 49 | No TBI vs. TBI + 50mM MPEP | 61.70 | 161.6 | -99.93 | 51 | 28 | 7.203 |
| 50 | No TBI vs. TBI + 100mM MPEP | 61.70 | 153.7 | -91.96 | 51 | 16 | 5.441 |
| 51 | No TBI vs. 50mM MPEP | 61.70 | 72.08 | -10.39 | 51 | 6 | 0.4080 |
| 52 | TBI vs. TBI + DMSO | 114.4 | 128.6 | -14.15 | 62 | 7 | 0.6017 |
| 53 | TBI vs. TBI + 3mM MPEP | 114.4 | 92.15 | 22.27 | 62 | 17 | 1.379 |
| 54 | TBI vs. TBI + 10mM MPEP | 114.4 | 130.5 | -16.03 | 62 | 10 | 0.7975 |
| 55 | TBI vs. TBI + 30mM MPEP | 114.4 | 133.0 | -18.58 | 62 | 25 | 1.330 |
| 56 | TBI vs. TBI + 50mM MPEP | 114.4 | 161.6 | -47.21 | 62 | 28 | 3.515 |
| 57 | TBI vs. TBI + 100mM MPEP | 114.4 | 153.7 | -39.24 | 62 | 16 | 2.372 |
| 58 | TBI vs. 50mM MPEP | 114.4 | 72.08 | 42.34 | 62 | 6 | 1.679 |
| 59 | TBI + DMSO vs. TBI + 3mM MPEP | 128.6 | 92.15 | 36.42 | 7 | 17 | 1.375 |
| 60 | TBI + DMSO vs. TBI + 10mM MPEP | 128.6 | 130.5 | -1.879 | 7 | 10 | 0.06463 |
| 61 | TBI + DMSO vs. TBI + 30mM MPEP | 128.6 | 133.0 | -4.429 | 7 | 25 | 0.1756 |
| 62 | TBI + DMSO vs. TBI + 50mM MPEP | 128.6 | 161.6 | -33.05 | 7 | 28 | 1.326 |

### Analysis associated with Figure 2E

| Kruskal-Wallis test<br>Multiple comparisons | | | | | | | |
|---|---|---|---|---|---|---|---|
| 63 | TBI + DMSO vs. TBI + 100mM MPEP | 128.6 | 153.7 | -25.08 | 7 | 16 | 0.9385 |
| 64 | TBI + DMSO vs. 50mM MPEP | 128.6 | 72.08 | 56.49 | 7 | 6 | 1.721 |
| 65 | TBI + 3mM MPEP vs. TBI + 10mM MPEP | 92.15 | 130.5 | -38.30 | 17 | 10 | 1.629 |
| 66 | TBI + 3mM MPEP vs. TBI + 30mM MPEP | 92.15 | 133.0 | -40.85 | 17 | 25 | 2.203 |
| 67 | TBI + 3mM MPEP vs. TBI + 50mM MPEP | 92.15 | 161.6 | -69.48 | 17 | 28 | 3.831 |
| 68 | TBI + 3mM MPEP vs. TBI + 100mM MPEP | 92.15 | 153.7 | -61.51 | 17 | 16 | 2.994 |
| 69 | TBI + 3mM MPEP vs. 50mM MPEP | 92.15 | 72.08 | 20.06 | 17 | 6 | 0.7163 |
| 70 | TBI + 10mM MPEP vs. TBI + 30mM MPEP | 130.5 | 133.0 | -2.550 | 10 | 25 | 0.1155 |
| 71 | TBI + 10mM MPEP vs. TBI + 50mM MPEP | 130.5 | 161.6 | -31.18 | 10 | 28 | 1.435 |
| 72 | TBI + 10mM MPEP vs. TBI + 100mM MPEP | 130.5 | 153.7 | -23.21 | 10 | 16 | 0.9760 |
| 73 | TBI + 10mM MPEP vs. 50mM MPEP | 130.5 | 72.08 | 58.37 | 10 | 6 | 1.916 |
| 74 | TBI + 30mM MPEP vs. TBI + 50mM MPEP | 133.0 | 161.6 | -28.63 | 25 | 28 | 1.764 |
| 75 | TBI + 30mM MPEP vs. TBI + 100mM MPEP | 133.0 | 153.7 | -20.66 | 25 | 16 | 1.094 |
| 76 | TBI + 30mM MPEP vs. 50mM MPEP | 133.0 | 72.08 | 60.92 | 25 | 6 | 2.272 |
| 77 | TBI + 50mM MPEP vs. TBI + 100mM MPEP | 161.6 | 153.7 | 7.969 | 28 | 16 | 0.4311 |
| 78 | TBI + 50mM MPEP vs. 50mM MPEP | 161.6 | 72.08 | 89.54 | 28 | 6 | 3.374 |
| 79 | TBI + 100mM MPEP vs. 50mM MPEP | 153.7 | 72.08 | 81.57 | 16 | 6 | 2.889 |

### Analysis associated with Figure 2F

| | **Kruskal-Wallis test**<br>ANOVA results | |
|---|---|---|
| 1 | Table Analyzed | chpg |
| 2 | | |
| 3 | **Kruskal-Wallis test** | |
| 4 | P value | 0.0342 |
| 5 | Exact or approximate P value? | Approximate |
| 6 | P value summary | * |
| 7 | Do the medians vary signif. (P < 0.05)? | Yes |
| 8 | Number of groups | 7 |
| 9 | Kruskal-Wallis statistic | 13.62 |
| 10 | | |
| 11 | **Data summary** | |
| 12 | Number of treatments (columns) | 7 |
| 13 | Number of values (total) | 77 |

### Analysis associated with Figure 2F

| | **Kruskal-Wallis test**<br>Multiple comparisons | | | | | | |
|---|---|---|---|---|---|---|---|
| 1 | Number of families | 1 | | | | | |
| 2 | Number of comparisons per family | 21 | | | | | |
| 3 | Alpha | 0.05 | | | | | |
| 4 | | | | | | | |
| 5 | **Dunn's multiple comparisons test** | **Mean rank diff.** | **Significant?** | **Summary** | **Adjusted P Value** | | |
| 6 | no TBI vs. TBI | -22.67 | No | ns | 0.0725 | A-B | |
| 7 | no TBI vs. 10μM CHPG + TBI | -11.70 | No | ns | >0.9999 | A-C | |
| 8 | no TBI vs. 30μM CHPG + TBI | -6.556 | No | ns | >0.9999 | A-D | |
| 9 | no TBI vs. 50μM CHPG + TBI | -4.350 | No | ns | >0.9999 | A-E | |
| 10 | no TBI vs. 100μM CHPG + TBI | -2.026 | No | ns | >0.9999 | A-F | |
| 11 | no TBI vs. 50μM CHPG no TBI | 0.000 | No | ns | >0.9999 | A-G | |
| 12 | TBI vs. 10μM CHPG + TBI | 10.97 | No | ns | >0.9999 | B-C | |
| 13 | TBI vs. 30μM CHPG + TBI | 16.11 | No | ns | 0.8984 | B-D | |
| 14 | TBI vs. 50μM CHPG + TBI | 18.32 | No | ns | 0.3806 | B-E | |
| 15 | TBI vs. 100μM CHPG + TBI | 20.64 | No | ns | 0.0525 | B-F | |
| 16 | TBI vs. 50μM CHPG no TBI | 22.67 | No | ns | 0.0725 | B-G | |
| 17 | 10μM CHPG + TBI vs. 30μM CHPG + TBI | 5.144 | No | ns | >0.9999 | C-D | |
| 18 | 10μM CHPG + TBI vs. 50μM CHPG + TBI | 7.350 | No | ns | >0.9999 | C-E | |
| 19 | 10μM CHPG + TBI vs. 100μM CHPG + TBI | 9.674 | No | ns | >0.9999 | C-F | |
| 20 | 10μM CHPG + TBI vs. 50μM CHPG no TBI | 11.70 | No | ns | >0.9999 | C-G | |
| 21 | 30μM CHPG + TBI vs. 50μM CHPG + TBI | 2.206 | No | ns | >0.9999 | D-E | |
| 22 | 30μM CHPG + TBI vs. 100μM CHPG + TBI | 4.529 | No | ns | >0.9999 | D-F | |
| 23 | 30μM CHPG + TBI vs. 50μM CHPG no TBI | 6.556 | No | ns | >0.9999 | D-G | |
| 24 | 50μM CHPG + TBI vs. 100μM CHPG + TBI | 2.324 | No | ns | >0.9999 | E-F | |
| 25 | 50μM CHPG + TBI vs. 50μM CHPG no TBI | 4.350 | No | ns | >0.9999 | E-G | |
| 26 | 100μM CHPG + TBI vs. 50μM CHPG no TBI | 2.026 | No | ns | >0.9999 | F-G | |
| 27 | | | | | | | |
| 28 | **Test details** | **Mean rank 1** | **Mean rank 2** | **Mean rank diff.** | **n1** | **n2** | **Z** |
| 29 | no TBI vs. TBI | 33.00 | 55.67 | -22.67 | 10 | 9 | 2.924 |
| 30 | no TBI vs. 10μM CHPG + TBI | 33.00 | 44.70 | -11.70 | 10 | 10 | 1.551 |
| 31 | no TBI vs. 30μM CHPG + TBI | 33.00 | 39.56 | -6.556 | 10 | 9 | 0.8457 |

### Analysis associated with Figure 2F

| | **Kruskal-Wallis test** Multiple comparisons | | | | | | |
|---|---|---|---|---|---|---|---|
| 32 | no TBI vs. 50μM CHPG + TBI | 33.00 | 37.35 | -4.350 | 10 | 10 | 0.5766 |
| 33 | no TBI vs. 100μM CHPG + TBI | 33.00 | 35.03 | -2.026 | 10 | 19 | 0.3074 |
| 34 | no TBI vs. 50μM CHPG no TBI | 33.00 | 33.00 | 0.000 | 10 | 10 | 0.000 |
| 35 | TBI vs. 10μM CHPG + TBI | 55.67 | 44.70 | 10.97 | 9 | 10 | 1.415 |
| 36 | TBI vs. 30μM CHPG + TBI | 55.67 | 39.56 | 16.11 | 9 | 9 | 2.026 |
| 37 | TBI vs. 50μM CHPG + TBI | 55.67 | 37.35 | 18.32 | 9 | 10 | 2.363 |
| 38 | TBI vs. 100μM CHPG + TBI | 55.67 | 35.03 | 20.64 | 9 | 19 | 3.024 |
| 39 | TBI vs. 50μM CHPG no TBI | 55.67 | 33.00 | 22.67 | 9 | 10 | 2.924 |
| 40 | 10μM CHPG + TBI vs. 30μM CHPG + TBI | 44.70 | 39.56 | 5.144 | 10 | 9 | 0.6637 |
| 41 | 10μM CHPG + TBI vs. 50μM CHPG + TBI | 44.70 | 37.35 | 7.350 | 10 | 10 | 0.9742 |
| 42 | 10μM CHPG + TBI vs. 100μM CHPG + TBI | 44.70 | 35.03 | 9.674 | 10 | 19 | 1.468 |
| 43 | 10μM CHPG + TBI vs. 50μM CHPG no TBI | 44.70 | 33.00 | 11.70 | 10 | 10 | 1.551 |
| 44 | 30μM CHPG + TBI vs. 50μM CHPG + TBI | 39.56 | 37.35 | 2.206 | 9 | 10 | 0.2845 |
| 45 | 30μM CHPG + TBI vs. 100μM CHPG + TBI | 39.56 | 35.03 | 4.529 | 9 | 19 | 0.6635 |
| 46 | 30μM CHPG + TBI vs. 50μM CHPG no TBI | 39.56 | 33.00 | 6.556 | 9 | 10 | 0.8457 |
| 47 | 50μM CHPG + TBI vs. 100μM CHPG + TBI | 37.35 | 35.03 | 2.324 | 10 | 19 | 0.3526 |
| 48 | 50μM CHPG + TBI vs. 50μM CHPG no TBI | 37.35 | 33.00 | 4.350 | 10 | 10 | 0.5766 |
| 49 | 100μM CHPG + TBI vs. 50μM CHPG no TBI | 35.03 | 33.00 | 2.026 | 19 | 10 | 0.3074 |
| 50 | | | | | | | |
| 51 | **Compact letter display** | | | | | | |
| 52 | TBI | A | | | | | |
| 53 | 10μM CHPG + TBI | A | | | | | |
| 54 | 30μM CHPG + TBI | A | | | | | |
| 55 | 50μM CHPG + TBI | A | | | | | |
| 56 | 100μM CHPG + TBI | A | | | | | |
| 57 | no TBI | A | | | | | |
| 58 | 50μM CHPG no TBI | A | | | | | |

### Analysis associated with Figure 2G

| | Kruskal-Wallis test<br>ANOVA results | |
|---|---|---|
| 1 | Table Analyzed | SCA_TBI_CHPG_MP |
| 2 | | |
| 3 | **Kruskal-Wallis test** | |
| 4 | P value | 0.0002 |
| 5 | Exact or approximate P value? | Approximate |
| 6 | P value summary | *** |
| 7 | Do the medians vary signif. (P < 0.05)? | Yes |
| 8 | Number of groups | 7 |
| 9 | Kruskal-Wallis statistic | 26.33 |
| 10 | | |
| 11 | **Data summary** | |
| 12 | Number of treatments (columns) | 7 |
| 13 | Number of values (total) | 106 |

### Analysis associated with Figure 2G

| | **Kruskal-Wallis test**<br>Multiple comparisons | | | | | | |
|---|---|---|---|---|---|---|---|
| 1 | Number of families | 1 | | | | | |
| 2 | Number of comparisons per family | 21 | | | | | |
| 3 | Alpha | 0.05 | | | | | |
| 4 | | | | | | | |
| 5 | **Dunn's multiple comparisons test** | **Mean rank diff.** | **Significant?** | **Summary** | **Adjusted P Value** | | |
| 6 | no TBI vs. TBI | -27.33 | Yes | * | 0.0191 | A-B | |
| 7 | no TBI vs. 3mM MPEP + 50mM CHPG + TBI | -8.250 | No | ns | >0.9999 | A-C | |
| 8 | no TBI vs. 10mM MPEP + 50mM CHPG + TBI | -20.58 | No | ns | 0.2186 | A-D | |
| 9 | no TBI vs. 30mM MPEP + 50mM CHPG + TBI | -40.55 | Yes | *** | 0.0004 | A-E | |
| 10 | no TBI vs. 50mM MPEP + 50mM CHPG + TBI | -6.188 | No | ns | >0.9999 | A-F | |
| 11 | no TBI vs. 30mM MPEP + 50mM CHPG no TBI | -11.42 | No | ns | >0.9999 | A-G | |
| 12 | TBI vs. 3mM MPEP + 50mM CHPG + TBI | 19.08 | No | ns | 0.4318 | B-C | |
| 13 | TBI vs. 10mM MPEP + 50mM CHPG + TBI | 6.758 | No | ns | >0.9999 | B-D | |
| 14 | TBI vs. 30mM MPEP + 50mM CHPG + TBI | -13.21 | No | ns | >0.9999 | B-E | |
| 15 | TBI vs. 50mM MPEP + 50mM CHPG + TBI | 21.15 | No | ns | 0.9261 | B-F | |
| 16 | TBI vs. 30mM MPEP + 50mM CHPG no TBI | 15.91 | No | ns | >0.9999 | B-G | |
| 17 | 3mM MPEP + 50mM CHPG + TBI vs. 10mM MPEP + 50mM CHPG + TBI | -12.33 | No | ns | >0.9999 | C-D | |
| 18 | 3mM MPEP + 50mM CHPG + TBI vs. 30mM MPEP + 50mM CHPG + TBI | -32.30 | Yes | * | 0.0135 | C-E | |
| 19 | 3mM MPEP + 50mM CHPG + TBI vs. 50mM MPEP + 50mM CHPG + TBI | 2.063 | No | ns | >0.9999 | C-F | |
| 20 | 3mM MPEP + 50mM CHPG + TBI vs. 30mM MPEP + 50mM CHPG no TBI | -3.173 | No | ns | >0.9999 | C-G | |
| 21 | 10mM MPEP + 50mM CHPG + TBI vs. 30mM MPEP + 50mM CHPG + TBI | -19.97 | No | ns | 0.6591 | D-E | |
| 22 | 10mM MPEP + 50mM CHPG + TBI vs. 50mM MPEP + 50mM CHPG + TBI | 14.39 | No | ns | >0.9999 | D-F | |
| 23 | 10mM MPEP + 50mM CHPG + TBI vs. 30mM MPEP + 50mM CHPG no TBI | 9.152 | No | ns | >0.9999 | D-G | |
| 24 | 30mM MPEP + 50mM CHPG + TBI vs. 50mM MPEP + 50mM CHPG + TBI | 34.36 | No | ns | 0.0584 | E-F | |
| 25 | 30mM MPEP + 50mM CHPG + TBI vs. 30mM MPEP + 50mM CHPG no TBI | 29.12 | No | ns | 0.0847 | E-G | |
| 26 | 50mM MPEP + 50mM CHPG + TBI vs. 30mM MPEP + 50mM CHPG no TBI | -5.236 | No | ns | >0.9999 | F-G | |
| 27 | | | | | | | |
| 28 | **Test details** | **Mean rank 1** | **Mean rank 2** | **Mean rank diff.** | **n1** | **n2** | **Z** |
| 29 | no TBI vs. TBI | 37.50 | 64.83 | -27.33 | 18 | 18 | 3.317 |
| 30 | no TBI vs. 3mM MPEP + 50mM CHPG + TBI | 37.50 | 45.75 | -8.250 | 18 | 18 | 1.001 |
| 31 | no TBI vs. 10mM MPEP + 50mM CHPG + TBI | 37.50 | 58.08 | -20.58 | 18 | 20 | 2.562 |

### Analysis associated with Figure 2G

| Kruskal-Wallis test<br>Multiple comparisons | | | | | | | |
|---|---|---|---|---|---|---|---|
| 32 | no TBI vs. 30mM MPEP + 50mM CHPG + TBI | 37.50 | 78.05 | -40.55 | 18 | 11 | 4.286 |
| 33 | no TBI vs. 50mM MPEP + 50mM CHPG + TBI | 37.50 | 43.69 | -6.188 | 18 | 8 | 0.5891 |
| 34 | no TBI vs. 30mM MPEP + 50mM CHPG no TBI | 37.50 | 48.92 | -11.42 | 18 | 13 | 1.270 |
| 35 | TBI vs. 3mM MPEP + 50mM CHPG + TBI | 64.83 | 45.75 | 19.08 | 18 | 18 | 2.316 |
| 36 | TBI vs. 10mM MPEP + 50mM CHPG + TBI | 64.83 | 58.08 | 6.758 | 18 | 20 | 0.8415 |
| 37 | TBI vs. 30mM MPEP + 50mM CHPG + TBI | 64.83 | 78.05 | -13.21 | 18 | 11 | 1.397 |
| 38 | TBI vs. 50mM MPEP + 50mM CHPG + TBI | 64.83 | 43.69 | 21.15 | 18 | 8 | 2.013 |
| 39 | TBI vs. 30mM MPEP + 50mM CHPG no TBI | 64.83 | 48.92 | 15.91 | 18 | 13 | 1.768 |
| 40 | 3mM MPEP + 50mM CHPG + TBI vs. 10mM MPEP + 50mM CHPG + TBI | 45.75 | 58.08 | -12.33 | 18 | 20 | 1.535 |
| 41 | 3mM MPEP + 50mM CHPG + TBI vs. 30mM MPEP + 50mM CHPG + TBI | 45.75 | 78.05 | -32.30 | 18 | 11 | 3.414 |
| 42 | 3mM MPEP + 50mM CHPG + TBI vs. 50mM MPEP + 50mM CHPG + TBI | 45.75 | 43.69 | 2.063 | 18 | 8 | 0.1964 |
| 43 | 3mM MPEP + 50mM CHPG + TBI vs. 30mM MPEP + 50mM CHPG no TBI | 45.75 | 48.92 | -3.173 | 18 | 13 | 0.3527 |
| 44 | 10mM MPEP + 50mM CHPG + TBI vs. 30mM MPEP + 50mM CHPG + TBI | 58.08 | 78.05 | -19.97 | 20 | 11 | 2.152 |
| 45 | 10mM MPEP + 50mM CHPG + TBI vs. 50mM MPEP + 50mM CHPG + TBI | 58.08 | 43.69 | 14.39 | 20 | 8 | 1.391 |
| 46 | 10mM MPEP + 50mM CHPG + TBI vs. 30mM MPEP + 50mM CHPG no TBI | 58.08 | 48.92 | 9.152 | 20 | 13 | 1.039 |
| 47 | 30mM MPEP + 50mM CHPG + TBI vs. 50mM MPEP + 50mM CHPG + TBI | 78.05 | 43.69 | 34.36 | 11 | 8 | 2.991 |
| 48 | 30mM MPEP + 50mM CHPG + TBI vs. 30mM MPEP + 50mM CHPG no TBI | 78.05 | 48.92 | 29.12 | 11 | 13 | 2.876 |
| 49 | 50mM MPEP + 50mM CHPG + TBI vs. 30mM MPEP + 50mM CHPG no TBI | 43.69 | 48.92 | -5.236 | 8 | 13 | 0.4713 |

### Analysis associated with Figure 3E

| | **Ordinary one-way ANOVA**<br>ANOVA results | | | | | |
|---|---|---|---|---|---|---|
| 1 | Table Analyzed | Data 1 | | | | |
| 2 | Data sets analyzed | A-F | | | | |
| 3 | | | | | | |
| 4 | **ANOVA summary** | | | | | |
| 5 | F | 55.30 | | | | |
| 6 | P value | <0.0001 | | | | |
| 7 | P value summary | **** | | | | |
| 8 | Significant diff. among means (P < 0.05)? | Yes | | | | |
| 9 | R squared | 0.8820 | | | | |
| 10 | | | | | | |
| 11 | **Brown-Forsythe test** | | | | | |
| 12 | F (DFn, DFd) | 0.9920 (5, 37) | | | | |
| 13 | P value | 0.4359 | | | | |
| 14 | P value summary | ns | | | | |
| 15 | Are SDs significantly different (P < 0.05)? | No | | | | |
| 16 | | | | | | |
| 17 | **Bartlett's test** | | | | | |
| 18 | Bartlett's statistic (corrected) | 7.104 | | | | |
| 19 | P value | 0.2130 | | | | |
| 20 | P value summary | ns | | | | |
| 21 | Are SDs significantly different (P < 0.05)? | No | | | | |
| 22 | | | | | | |
| 23 | **ANOVA table** | **SS** | **DF** | **MS** | **F (DFn, DFd)** | **P value** |
| 24 | Treatment (between columns) | 9486 | 5 | 1897 | F (5, 37) = 55.30 | P<0.0001 |
| 25 | Residual (within columns) | 1269 | 37 | 34.30 | | |
| 26 | Total | 10755 | 42 | | | |
| 27 | | | | | | |
| 28 | **Data summary** | | | | | |
| 29 | Number of treatments (columns) | 6 | | | | |
| 30 | Number of values (total) | 43 | | | | |

### Analysis associated with Figure 3E

| | **Ordinary one-way ANOVA** Multiple comparisons | | | | | | | | |
|---|---|---|---|---|---|---|---|---|---|
| 1 | Number of families | 1 | | | | | | | |
| 2 | Number of comparisons per family | 15 | | | | | | | |
| 3 | Alpha | 0.05 | | | | | | | |
| 4 | | | | | | | | | |
| 5 | **Tukey's multiple comparisons test** | **Mean Diff.** | **95.00% CI of diff.** | **Below threshold?** | **Summary** | **Adjusted P Value** | | | |
| 6 | no tbi vs. tbi | -14.07 | -23.18 to -4.965 | Yes | *** | 0.0006 | A-B | | |
| 7 | no tbi vs. tbi mpep 50µM | -38.35 | -47.22 to -29.48 | Yes | **** | <0.0001 | A-C | | |
| 8 | no tbi vs. tbi chpg 50µM | -1.905 | -10.77 to 6.962 | No | ns | 0.9866 | A-D | | |
| 9 | no tbi vs. mpep 50µM | -14.37 | -24.67 to -4.069 | Yes | ** | 0.0021 | A-E | | |
| 10 | no tbi vs. chpg 50µM | 3.829 | -6.474 to 14.13 | No | ns | 0.8715 | A-F | | |
| 11 | tbi vs. tbi mpep 50µM | -24.28 | -32.83 to -15.73 | Yes | **** | <0.0001 | B-C | | |
| 12 | tbi vs. tbi chpg 50µM | 12.17 | 3.617 to 20.72 | Yes | ** | 0.0017 | B-D | | |
| 13 | tbi vs. mpep 50µM | -0.3000 | -10.33 to 9.731 | No | ns | >0.9999 | B-E | | |
| 14 | tbi vs. chpg 50µM | 17.90 | 7.869 to 27.93 | Yes | **** | <0.0001 | B-F | | |
| 15 | tbi mpep 50µM vs. tbi chpg 50µM | 36.44 | 28.15 to 44.74 | Yes | **** | <0.0001 | C-D | | |
| 16 | tbi mpep 50µM vs. mpep 50µM | 23.98 | 14.16 to 33.79 | Yes | **** | <0.0001 | C-E | | |
| 17 | tbi mpep 50µM vs. chpg 50µM | 42.18 | 32.36 to 51.99 | Yes | **** | <0.0001 | C-F | | |
| 18 | tbi chpg 50µM vs. mpep 50µM | -12.47 | -22.28 to -2.652 | Yes | ** | 0.0062 | D-E | | |
| 19 | tbi chpg 50µM vs. chpg 50µM | 5.733 | -4.081 to 15.55 | No | ns | 0.5060 | D-F | | |
| 20 | mpep 50µM vs. chpg 50µM | 18.20 | 7.072 to 29.33 | Yes | *** | 0.0003 | E-F | | |
| 21 | | | | | | | | | |
| 22 | **Test details** | **Mean 1** | **Mean 2** | **Mean Diff.** | **SE of diff.** | **n1** | **n2** | **q** | **DF** |
| 23 | no tbi vs. tbi | 11.43 | 25.50 | -14.07 | 3.031 | 7 | 8 | 6.565 | 37 |
| 24 | no tbi vs. tbi mpep 50µM | 11.43 | 49.78 | -38.35 | 2.952 | 7 | 9 | 18.37 | 37 |
| 25 | no tbi vs. tbi chpg 50µM | 11.43 | 13.33 | -1.905 | 2.952 | 7 | 9 | 0.9126 | 37 |
| 26 | no tbi vs. mpep 50µM | 11.43 | 25.80 | -14.37 | 3.430 | 7 | 5 | 5.926 | 37 |
| 27 | no tbi vs. chpg 50µM | 11.43 | 7.600 | 3.829 | 3.430 | 7 | 5 | 1.579 | 37 |
| 28 | tbi vs. tbi mpep 50µM | 25.50 | 49.78 | -24.28 | 2.846 | 8 | 9 | 12.06 | 37 |
| 29 | tbi vs. tbi chpg 50µM | 25.50 | 13.33 | 12.17 | 2.846 | 8 | 9 | 6.046 | 37 |
| 30 | tbi vs. mpep 50µM | 25.50 | 25.80 | -0.3000 | 3.339 | 8 | 5 | 0.1271 | 37 |
| 31 | tbi vs. chpg 50µM | 25.50 | 7.600 | 17.90 | 3.339 | 8 | 5 | 7.581 | 37 |

### Analysis associated with Figure 3E

| | **Ordinary one-way ANOVA** Multiple comparisons | | | | | | | | |
|---|---|---|---|---|---|---|---|---|---|
| 32 | tbi mpep 50µM vs. tbi chpg 50µM | 49.78 | 13.33 | 36.44 | 2.761 | 9 | 9 | 18.67 | 37 |
| 33 | tbi mpep 50µM vs. mpep 50µM | 49.78 | 25.80 | 23.98 | 3.267 | 9 | 5 | 10.38 | 37 |
| 34 | tbi mpep 50µM vs. chpg 50µM | 49.78 | 7.600 | 42.18 | 3.267 | 9 | 5 | 18.26 | 37 |
| 35 | tbi chpg 50µM vs. mpep 50µM | 13.33 | 25.80 | -12.47 | 3.267 | 9 | 5 | 5.397 | 37 |
| 36 | tbi chpg 50µM vs. chpg 50µM | 13.33 | 7.600 | 5.733 | 3.267 | 9 | 5 | 2.482 | 37 |
| 37 | mpep 50µM vs. chpg 50µM | 25.80 | 7.600 | 18.20 | 3.704 | 5 | 5 | 6.948 | 37 |
| 38 | | | | | | | | | |
| 39 | **Compact letter display** | | | | | | | | |
| 40 | tbi mpep 50µM | A | | | | | | | |
| 41 | mpep 50µM | B | | | | | | | |
| 42 | tbi | B | | | | | | | |
| 43 | tbi chpg 50µM | C | | | | | | | |
| 44 | no tbi | C | | | | | | | |
| 45 | chpg 50µM | C | | | | | | | |

### Analysis associated with Figure 4C

| | **Kruskal-Wallis test**<br>ANOVA results | |
|---|---|---|
| 1 | Table Analyzed | Data 1 |
| 2 | | |
| 3 | **Kruskal-Wallis test** | |
| 4 | P value | <0.0001 |
| 5 | Exact or approximate P value? | Approximate |
| 6 | P value summary | **** |
| 7 | Do the medians vary signif. (P < 0.05)? | Yes |
| 8 | Number of groups | 6 |
| 9 | Kruskal-Wallis statistic | 32.35 |
| 10 | | |
| 11 | **Data summary** | |
| 12 | Number of treatments (columns) | 6 |
| 13 | Number of values (total) | 92 |

### Analysis associated with Figure 4C

| | Kruskal-Wallis test<br>Multiple comparisons | | | | | | |
|---|---|---|---|---|---|---|---|
| 1 | Number of families | 1 | | | | | |
| 2 | Number of comparisons per family | 15 | | | | | |
| 3 | Alpha | 0.05 | | | | | |
| 4 | | | | | | | |
| 5 | **Dunn's multiple comparisons test** | **Mean rank diff.** | **Significant?** | **Summary** | **Adjusted P Value** | | |
| 6 | PTZ (5mM) vs. PTZ (5mM) + MPEP (50μM) | -15.87 | No | ns | 0.1449 | A-B | |
| 7 | PTZ (5mM) vs. PTZ (5mM) + CHPG (50μM) | 16.50 | No | ns | 0.2473 | A-C | |
| 8 | PTZ (5mM) vs. MPEP (50μM) | 6.500 | No | ns | >0.9999 | A-D | |
| 9 | PTZ (5mM) vs. CHPG (50μM) | 16.50 | No | ns | 0.9522 | A-E | |
| 10 | PTZ (5mM) vs. No treatment | 16.50 | No | ns | 0.9522 | A-F | |
| 11 | PTZ (5mM) + MPEP (50μM) vs. PTZ (5mM) + CHPG (50μM) | 32.37 | Yes | **** | <0.0001 | B-C | |
| 12 | PTZ (5mM) + MPEP (50μM) vs. MPEP (50μM) | 22.37 | No | ns | 0.1724 | B-D | |
| 13 | PTZ (5mM) + MPEP (50μM) vs. CHPG (50μM) | 32.37 | Yes | ** | 0.0038 | B-E | |
| 14 | PTZ (5mM) + MPEP (50μM) vs. No treatment | 32.37 | Yes | ** | 0.0038 | B-F | |
| 15 | PTZ (5mM) + CHPG (50μM) vs. MPEP (50μM) | -10.00 | No | ns | >0.9999 | C-D | |
| 16 | PTZ (5mM) + CHPG (50μM) vs. CHPG (50μM) | 0.000 | No | ns | >0.9999 | C-E | |
| 17 | PTZ (5mM) + CHPG (50μM) vs. No treatment | 0.000 | No | ns | >0.9999 | C-F | |
| 18 | MPEP (50μM) vs. CHPG (50μM) | 10.00 | No | ns | >0.9999 | D-E | |
| 19 | MPEP (50μM) vs. No treatment | 10.00 | No | ns | >0.9999 | D-F | |
| 20 | CHPG (50μM) vs. No treatment | 0.000 | No | ns | >0.9999 | E-F | |
| 21 | | | | | | | |
| 22 | **Test details** | **Mean rank 1** | **Mean rank 2** | **Mean rank diff.** | **n1** | **n2** | **Z** |
| 23 | PTZ (5mM) vs. PTZ (5mM) + MPEP (50μM) | 48.50 | 64.37 | -15.87 | 25 | 26 | 2.588 |
| 24 | PTZ (5mM) vs. PTZ (5mM) + CHPG (50μM) | 48.50 | 32.00 | 16.50 | 25 | 17 | 2.398 |
| 25 | PTZ (5mM) vs. MPEP (50μM) | 48.50 | 42.00 | 6.500 | 25 | 8 | 0.7311 |
| 26 | PTZ (5mM) vs. CHPG (50μM) | 48.50 | 32.00 | 16.50 | 25 | 8 | 1.856 |
| 27 | PTZ (5mM) vs. No treatment | 48.50 | 32.00 | 16.50 | 25 | 8 | 1.856 |
| 28 | PTZ (5mM) + MPEP (50μM) vs. PTZ (5mM) + CHPG (50μM) | 64.37 | 32.00 | 32.37 | 26 | 17 | 4.741 |
| 29 | PTZ (5mM) + MPEP (50μM) vs. MPEP (50μM) | 64.37 | 42.00 | 22.37 | 26 | 8 | 2.527 |
| 30 | PTZ (5mM) + MPEP (50μM) vs. CHPG (50μM) | 64.37 | 32.00 | 32.37 | 26 | 8 | 3.657 |
| 31 | PTZ (5mM) + MPEP (50μM) vs. No treatment | 64.37 | 32.00 | 32.37 | 26 | 8 | 3.657 |

### Analysis associated with Figure 4C

| Kruskal-Wallis test<br>Multiple comparisons | | | | | | | |
|---|---|---|---|---|---|---|---|
| 32 | PTZ (5mM) + CHPG (50µM) vs. MPEP (50µM) | 32.00 | 42.00 | -10.00 | 17 | 8 | 1.066 |
| 33 | PTZ (5mM) + CHPG (50µM) vs. CHPG (50µM) | 32.00 | 32.00 | 0.000 | 17 | 8 | 0.000 |
| 34 | PTZ (5mM) + CHPG (50µM) vs. No treatment | 32.00 | 32.00 | 0.000 | 17 | 8 | 0.000 |
| 35 | MPEP (50µM) vs. CHPG (50µM) | 42.00 | 32.00 | 10.00 | 8 | 8 | 0.9137 |
| 36 | MPEP (50µM) vs. No treatment | 42.00 | 32.00 | 10.00 | 8 | 8 | 0.9137 |
| 37 | CHPG (50µM) vs. No treatment | 32.00 | 32.00 | 0.000 | 8 | 8 | 0.000 |
| 38 | | | | | | | |
| 39 | **Compact letter display** | | | | | | |
| 40 | PTZ (5mM) + MPEP (50µM) | A | | | | | |
| 41 | PTZ (5mM) | AB | | | | | |
| 42 | MPEP (50µM) | AB | | | | | |
| 43 | PTZ (5mM) + CHPG (50µM) | B | | | | | |
| 44 | CHPG (50µM) | B | | | | | |
| 45 | No treatment | B | | | | | |

### Analysis associated with Figure 5C

| | **Kruskal-Wallis test**<br>ANOVA results | |
|---|---|---|
| 1 | Table Analyzed | Data 2 |
| 2 | | |
| 3 | **Kruskal-Wallis test** | |
| 4 | P value | <0.0001 |
| 5 | Exact or approximate P value? | Approximate |
| 6 | P value summary | **** |
| 7 | Do the medians vary signif. (P < 0.05)? | Yes |
| 8 | Number of groups | 7 |
| 9 | Kruskal-Wallis statistic | 30.67 |
| 10 | | |
| 11 | **Data summary** | |
| 12 | Number of treatments (columns) | 7 |
| 13 | Number of values (total) | 147 |

### Analysis associated with Figure 5C

| | **Kruskal-Wallis test**<br>Multiple comparisons | | | | | | |
|---|---|---|---|---|---|---|---|
| 1 | Number of families | 1 | | | | | |
| 2 | Number of comparisons per family | 21 | | | | | |
| 3 | Alpha | 0.05 | | | | | |
| 4 | | | | | | | |
| 5 | **Dunn's multiple comparisons test** | **Mean rank diff.** | **Significant?** | **Summary** | **Adjusted P Value** | | |
| 6 | no TBI vs. TBI | -33.60 | No | ns | 0.2174 | A-B | |
| 7 | no TBI vs. TBI + 50µM CHPG | -1.825 | No | ns | >0.9999 | A-C | |
| 8 | no TBI vs. TBI + 10µM LY294 | -50.96 | Yes | ** | 0.0098 | A-D | |
| 9 | no TBI vs. TBI + 50µM CHPG + 5µM LY294 | -29.06 | No | ns | 0.2692 | A-E | |
| 10 | no TBI vs. TBI + 50µM CHPG + 10µM LY294 | -22.39 | No | ns | >0.9999 | A-F | |
| 11 | no TBI vs. TBI + 50µM CHPG + 15µM LY294 | -50.83 | Yes | ** | 0.0012 | A-G | |
| 12 | TBI vs. TBI + 50µM CHPG | 31.77 | No | ns | 0.2172 | B-C | |
| 13 | TBI vs. TBI + 10µM LY294 | -17.36 | No | ns | >0.9999 | B-D | |
| 14 | TBI vs. TBI + 50µM CHPG + 5µM LY294 | 4.535 | No | ns | >0.9999 | B-E | |
| 15 | TBI vs. TBI + 50µM CHPG + 10µM LY294 | 11.20 | No | ns | >0.9999 | B-F | |
| 16 | TBI vs. TBI + 50µM CHPG + 15µM LY294 | -17.23 | No | ns | >0.9999 | B-G | |
| 17 | TBI + 50µM CHPG vs. TBI + 10µM LY294 | -49.13 | Yes | ** | 0.0088 | C-D | |
| 18 | TBI + 50µM CHPG vs. TBI + 50µM CHPG + 5µM LY294 | -27.24 | No | ns | 0.2568 | C-E | |
| 19 | TBI + 50µM CHPG vs. TBI + 50µM CHPG + 10µM LY294 | -20.57 | No | ns | >0.9999 | C-F | |
| 20 | TBI + 50µM CHPG vs. TBI + 50µM CHPG + 15µM LY294 | -49.00 | Yes | *** | 0.0008 | C-G | |
| 21 | TBI + 10µM LY294 vs. TBI + 50µM CHPG + 5µM LY294 | 21.90 | No | ns | >0.9999 | D-E | |
| 22 | TBI + 10µM LY294 vs. TBI + 50µM CHPG + 10µM LY294 | 28.57 | No | ns | 0.6289 | D-F | |
| 23 | TBI + 10µM LY294 vs. TBI + 50µM CHPG + 15µM LY294 | 0.1310 | No | ns | >0.9999 | D-G | |
| 24 | TBI + 50µM CHPG + 5µM LY294 vs. TBI + 50µM CHPG + 10µM LY294 | 6.670 | No | ns | >0.9999 | E-F | |
| 25 | TBI + 50µM CHPG + 5µM LY294 vs. TBI + 50µM CHPG + 15µM LY294 | -21.76 | No | ns | 0.8851 | E-G | |
| 26 | TBI + 50µM CHPG + 10µM LY294 vs. TBI + 50µM CHPG + 15µM LY294 | -28.43 | No | ns | 0.2059 | F-G | |
| 27 | | | | | | | |
| 28 | **Test details** | **Mean rank 1** | **Mean rank 2** | **Mean rank diff.** | **n1** | **n2** | **Z** |
| 29 | no TBI vs. TBI | 47.63 | 81.22 | -33.60 | 16 | 18 | 2.564 |
| 30 | no TBI vs. TBI + 50µM CHPG | 47.63 | 49.45 | -1.825 | 16 | 20 | 0.1427 |
| 31 | no TBI vs. TBI + 10µM LY294 | 47.63 | 98.58 | -50.96 | 16 | 12 | 3.499 |

### Analysis associated with Figure 5C

| Kruskal-Wallis test<br>Multiple comparisons | | | | | | | |
|---|---|---|---|---|---|---|---|
| 32 | no TBI vs. TBI + 50µM CHPG + 5µM LY294 | 47.63 | 76.69 | -29.06 | 16 | 32 | 2.489 |
| 33 | no TBI vs. TBI + 50µM CHPG + 10µM LY294 | 47.63 | 70.02 | -22.39 | 16 | 28 | 1.873 |
| 34 | no TBI vs. TBI + 50µM CHPG + 15µM LY294 | 47.63 | 98.45 | -50.83 | 16 | 21 | 4.016 |
| 35 | TBI vs. TBI + 50µM CHPG | 81.22 | 49.45 | 31.77 | 18 | 20 | 2.564 |
| 36 | TBI vs. TBI + 10µM LY294 | 81.22 | 98.58 | -17.36 | 18 | 12 | 1.221 |
| 37 | TBI vs. TBI + 50µM CHPG + 5µM LY294 | 81.22 | 76.69 | 4.535 | 18 | 32 | 0.4036 |
| 38 | TBI vs. TBI + 50µM CHPG + 10µM LY294 | 81.22 | 70.02 | 11.20 | 18 | 28 | 0.9724 |
| 39 | TBI vs. TBI + 50µM CHPG + 15µM LY294 | 81.22 | 98.45 | -17.23 | 18 | 21 | 1.406 |
| 40 | TBI + 50µM CHPG vs. TBI + 10µM LY294 | 49.45 | 98.58 | -49.13 | 20 | 12 | 3.528 |
| 41 | TBI + 50µM CHPG vs. TBI + 50µM CHPG + 5µM LY294 | 49.45 | 76.69 | -27.24 | 20 | 32 | 2.505 |
| 42 | TBI + 50µM CHPG vs. TBI + 50µM CHPG + 10µM LY294 | 49.45 | 70.02 | -20.57 | 20 | 28 | 1.842 |
| 43 | TBI + 50µM CHPG vs. TBI + 50µM CHPG + 15µM LY294 | 49.45 | 98.45 | -49.00 | 20 | 21 | 4.112 |
| 44 | TBI + 10µM LY294 vs. TBI + 50µM CHPG + 5µM LY294 | 98.58 | 76.69 | 21.90 | 12 | 32 | 1.696 |
| 45 | TBI + 10µM LY294 vs. TBI + 50µM CHPG + 10µM LY294 | 98.58 | 70.02 | 28.57 | 12 | 28 | 2.171 |
| 46 | TBI + 10µM LY294 vs. TBI + 50µM CHPG + 15µM LY294 | 98.58 | 98.45 | 0.1310 | 12 | 21 | 0.009488 |
| 47 | TBI + 50µM CHPG + 5µM LY294 vs. TBI + 50µM CHPG + 10µM LY294 | 76.69 | 70.02 | 6.670 | 32 | 28 | 0.6758 |
| 48 | TBI + 50µM CHPG + 5µM LY294 vs. TBI + 50µM CHPG + 15µM LY294 | 76.69 | 98.45 | -21.76 | 32 | 21 | 2.032 |
| 49 | TBI + 50µM CHPG + 10µM LY294 vs. TBI + 50µM CHPG + 15µM LY294 | 70.02 | 98.45 | -28.43 | 28 | 21 | 2.583 |

### Analysis associated with Figure 5D

| Mann-Whitney test | | |
|---|---|---|
| 1 | Table Analyzed | prp2;UA3171 20mL clamp 300g 3x |
| 2 | | |
| 3 | Column F | *prp2*$^{-/-}$ TBI |
| 4 | vs. | vs. |
| 5 | Column B | WT TBI |
| 6 | | |
| 7 | **Mann Whitney test** | |
| 8 | P value | 0.0081 |
| 9 | Exact or approximate P value? | Exact |
| 10 | P value summary | ** |
| 11 | Significantly different (P < 0.05)? | Yes |
| 12 | One- or two-tailed P value? | One-tailed |
| 13 | Sum of ranks in column B,F | 995.5 , 3191 |
| 14 | Mann-Whitney U | 617.5 |
| 15 | | |
| 16 | **Difference between medians** | |
| 17 | Median of column B | 0.000, n=27 |
| 18 | Median of column F | 1.000, n=64 |
| 19 | Difference: Actual | 1.000 |
| 20 | Difference: Hodges-Lehmann | 0.000 |

### Analysis associated with Figure 5E

| | **Kruskal-Wallis test**<br>ANOVA results | |
|---|---|---|
| 1 | Table Analyzed | prp2;UA3171 20mL clamp 300g 3x |
| 2 | | |
| 3 | **Kruskal-Wallis test** | |
| 4 | P value | <0.0001 |
| 5 | Exact or approximate P value? | Approximate |
| 6 | P value summary | **** |
| 7 | Do the medians vary signif. (P < 0.05)? | Yes |
| 8 | Number of groups | 3 |
| 9 | Kruskal-Wallis statistic | 24.07 |
| 10 | | |
| 11 | **Data summary** | |
| 12 | Number of treatments (columns) | 3 |
| 13 | Number of values (total) | 107 |

### Analysis associated with Figure 5E

| Kruskal-Wallis test<br>Multiple comparisons | | | | | | | |
|---|---|---|---|---|---|---|---|
| 1 | Number of families | 1 | | | | | |
| 2 | Number of comparisons per family | 3 | | | | | |
| 3 | Alpha | 0.05 | | | | | |
| 4 | | | | | | | |
| 5 | **Dunn's multiple comparisons test** | **Mean rank diff.** | **Significant?** | **Summary** | **Adjusted P V** | | |
| 6 | *prp2*$^{-/-}$no TBI vs. *prp2*$^{-/-}$ TBI | -25.58 | Yes | **** | <0.0001 | E-F | |
| 7 | *prp2*$^{-/-}$no TBI vs. *prp2*$^{-/-}$ TBI + 50µM CHPG | -1.655 | No | ns | >0.9999 | E-H | |
| 8 | *prp2*$^{-/-}$ TBI vs. *prp2*$^{-/-}$ TBI + 50µM CHPG | 23.93 | Yes | * | 0.0208 | F-H | |
| 9 | | | | | | | |
| 10 | **Test details** | **Mean rank 1** | **Mean rank 2** | **Mean rank d** | **n1** | **n2** | **Z** |
| 11 | *prp2*$^{-/-}$no TBI vs. *prp2*$^{-/-}$ TBI | 38.55 | 64.13 | -25.58 | 33 | 64 | 4.580 |
| 12 | *prp2*$^{-/-}$no TBI vs. *prp2*$^{-/-}$ TBI + 50µM CHPG | 38.55 | 40.20 | -1.655 | 33 | 10 | 0.1759 |
| 13 | *prp2*$^{-/-}$ TBI vs. *prp2*$^{-/-}$ TBI + 50µM CHPG | 64.13 | 40.20 | 23.93 | 64 | 10 | 2.700 |
| 14 | | | | | | | |
| 15 | **Compact letter display** | | | | | | |
| 16 | *prp2*$^{-/-}$ TBI | A | | | | | |
| 17 | *prp2*$^{-/-}$ TBI + 50µM CHPG | B | | | | | |
| 18 | *prp2*$^{-/-}$no TBI | B | | | | | |

### Analysis associated with Figure 6A

| Ordinary one-way ANOVA<br>ANOVA results | | | | | | |
|---|---|---|---|---|---|---|
| 1 | Table Analyzed | 6dpf seizure mean 300g | | | | |
| 2 | Data sets analyzed | A-D | | | | |
| 3 | | | | | | |
| 4 | **ANOVA summary** | | | | | |
| 5 | F | 9.028 | | | | |
| 6 | P value | <0.0001 | | | | |
| 7 | P value summary | **** | | | | |
| 8 | Significant diff. among means (P < 0.05)? | Yes | | | | |
| 9 | R squared | 0.1735 | | | | |
| 10 | | | | | | |
| 11 | **Brown-Forsythe test** | | | | | |
| 12 | F (DFn, DFd) | 6.697 (3, 129) | | | | |
| 13 | P value | 0.0003 | | | | |
| 14 | P value summary | *** | | | | |
| 15 | Are SDs significantly different (P < 0.05)? | Yes | | | | |
| 16 | | | | | | |
| 17 | **Bartlett's test** | | | | | |
| 18 | Bartlett's statistic (corrected) | 15.60 | | | | |
| 19 | P value | 0.0014 | | | | |
| 20 | P value summary | ** | | | | |
| 21 | Are SDs significantly different (P < 0.05)? | Yes | | | | |
| 22 | | | | | | |
| 23 | **ANOVA table** | **SS** | **DF** | **MS** | **F (DFn, DFd)** | **P value** |
| 24 | Treatment (between columns) | 0.02938 | 3 | 0.009795 | F (3, 129) = 9.028 | P<0.0001 |
| 25 | Residual (within columns) | 0.1400 | 129 | 0.001085 | | |
| 26 | Total | 0.1693 | 132 | | | |
| 27 | | | | | | |
| 28 | **Data summary** | | | | | |
| 29 | Number of treatments (columns) | 4 | | | | |
| 30 | Number of values (total) | 133 | | | | |

### Analysis associated with Figure 6A

| | **Ordinary one-way ANOVA**<br>Multiple comparisons | | | | | | | | |
|---|---|---|---|---|---|---|---|---|---|
| 1 | Number of families | 1 | | | | | | | |
| 2 | Number of comparisons per family | 6 | | | | | | | |
| 3 | Alpha | 0.05 | | | | | | | |
| 4 | | | | | | | | | |
| 5 | **Tukey's multiple comparisons test** | **Mean Diff.** | **95.00% CI of diff.** | **Below threshold?** | **Summary** | **Adjusted P Value** | | | |
| 6 | AB no TBI vs. AB TBI | 0.006304 | -0.01405 to 0.02666 | No | ns | 0.8514 | A-B | | |
| 7 | AB no TBI vs. *prp2*$^{-/-}$no TBI | -0.003365 | -0.02470 to 0.01797 | No | ns | 0.9766 | A-C | | |
| 8 | AB no TBI vs. *prp2*$^{-/-}$TBI | -0.03256 | -0.05353 to -0.01159 | Yes | *** | 0.0005 | A-D | | |
| 9 | AB TBI vs. *prp2*$^{-/-}$no TBI | -0.009669 | -0.03086 to 0.01153 | No | ns | 0.6359 | B-C | | |
| 10 | AB TBI vs. *prp2*$^{-/-}$TBI | -0.03886 | -0.05969 to -0.01803 | Yes | **** | <0.0001 | B-D | | |
| 11 | *prp2*$^{-/-}$no TBI vs. *prp2*$^{-/-}$TBI | -0.02919 | -0.05098 to -0.007405 | Yes | ** | 0.0037 | C-D | | |
| 12 | | | | | | | | | |
| 13 | **Test details** | **Mean 1** | **Mean 2** | **Mean Diff.** | **SE of diff.** | **n1** | **n2** | **q** | **DF** |
| 14 | AB no TBI vs. AB TBI | 0.04011 | 0.03381 | 0.006304 | 0.007819 | 35 | 36 | 1.140 | 129 |
| 15 | AB no TBI vs. *prp2*$^{-/-}$no TBI | 0.04011 | 0.04348 | -0.003365 | 0.008195 | 35 | 30 | 0.5806 | 129 |
| 16 | AB no TBI vs. *prp2*$^{-/-}$TBI | 0.04011 | 0.07267 | -0.03256 | 0.008056 | 35 | 32 | 5.715 | 129 |
| 17 | AB TBI vs. *prp2*$^{-/-}$no TBI | 0.03381 | 0.04348 | -0.009669 | 0.008143 | 36 | 30 | 1.679 | 129 |
| 18 | AB TBI vs. *prp2*$^{-/-}$TBI | 0.03381 | 0.07267 | -0.03886 | 0.008003 | 36 | 32 | 6.868 | 129 |
| 19 | *prp2*$^{-/-}$no TBI vs. *prp2*$^{-/-}$TBI | 0.04348 | 0.07267 | -0.02919 | 0.008371 | 30 | 32 | 4.932 | 129 |
| 20 | | | | | | | | | |
| 21 | **Compact letter display** | | | | | | | | |
| 22 | *prp2*$^{-/-}$TBI | A | | | | | | | |
| 23 | *prp2*$^{-/-}$no TBI | B | | | | | | | |
| 24 | AB no TBI | B | | | | | | | |
| 25 | AB TBI | B | | | | | | | |

### Analysis associated with Figure 6C

| Ordinary one-way ANOVA<br>ANOVA results | | | | | | |
|---|---|---|---|---|---|---|
| 1 | Table Analyzed | tbi + MPEP | | | | |
| 2 | Data sets analyzed | A-H | | | | |
| 3 | | | | | | |
| 4 | **ANOVA summary** | | | | | |
| 5 | F | 22.75 | | | | |
| 6 | P value | <0.0001 | | | | |
| 7 | P value summary | **** | | | | |
| 8 | Significant diff. among means (P < 0.05)? | Yes | | | | |
| 9 | R squared | 0.3537 | | | | |
| 10 | | | | | | |
| 11 | **Brown-Forsythe test** | | | | | |
| 12 | F (DFn, DFd) | 6.175 (7, 291) | | | | |
| 13 | P value | <0.0001 | | | | |
| 14 | P value summary | **** | | | | |
| 15 | Are SDs significantly different (P < 0.05)? | Yes | | | | |
| 16 | | | | | | |
| 17 | **Bartlett's test** | | | | | |
| 18 | Bartlett's statistic (corrected) | 62.24 | | | | |
| 19 | P value | <0.0001 | | | | |
| 20 | P value summary | **** | | | | |
| 21 | Are SDs significantly different (P < 0.05)? | Yes | | | | |
| 22 | | | | | | |
| 23 | **ANOVA table** | **SS** | **DF** | **MS** | **F (DFn, DFd)** | **P value** |
| 24 | Treatment (between columns) | 0.2656 | 7 | 0.03794 | F (7, 291) = 22.75 | P<0.0001 |
| 25 | Residual (within columns) | 0.4854 | 291 | 0.001668 | | |
| 26 | Total | 0.7510 | 298 | | | |
| 27 | | | | | | |
| 28 | **Data summary** | | | | | |
| 29 | Number of treatments (columns) | 8 | | | | |
| 30 | Number of values (total) | 299 | | | | |

### Analysis associated with Figure 6C

| Ordinary one-way ANOVA<br>Multiple comparisons | | | | | | | |
|---|---|---|---|---|---|---|---|
| 1 | Number of families | 1 | | | | | |
| 2 | Number of comparisons per family | 28 | | | | | |
| 3 | Alpha | 0.05 | | | | | |
| 4 | | | | | | | |
| 5 | **Tukey's multiple comparisons test** | **Mean Diff.** | **95.00% CI of diff.** | **Below thresh** | **Summary** | **Adjusted P V** | |
| 6 | AB no treatment vs. AB TBI | 0.003136 | -0.02013 to 0.02640 | No | ns | >0.9999 | A-B |
| 7 | AB no treatment vs. AB 10uM MPEP | -0.001805 | -0.03559 to 0.03198 | No | ns | >0.9999 | A-C |
| 8 | AB no treatment vs. AB 10uM MPEP + TBI | -0.02521 | -0.06056 to 0.01013 | No | ns | 0.3683 | A-D |
| 9 | AB no treatment vs. *prp2*$^{-/-}$ no treatment | 0.005504 | -0.01828 to 0.02929 | No | ns | 0.9968 | A-E |
| 10 | AB no treatment vs. *prp2*$^{-/-}$ + TBI | -0.02272 | -0.04571 to 0.0002655 | No | ns | 0.0553 | A-F |
| 11 | AB no treatment vs. *prp2*$^{-/-}$ 10uM MPEP | -0.01376 | -0.05001 to 0.02249 | No | ns | 0.9427 | A-G |
| 12 | AB no treatment vs. *prp2*$^{-/-}$ 10uM MPEP + TBI | -0.1186 | -0.1517 to -0.08547 | Yes | **** | <0.0001 | A-H |
| 13 | AB TBI vs. AB 10uM MPEP | -0.004941 | -0.03852 to 0.02864 | No | ns | 0.9998 | B-C |
| 14 | AB TBI vs. AB 10uM MPEP + TBI | -0.02835 | -0.06350 to 0.006798 | No | ns | 0.2159 | B-D |
| 15 | AB TBI vs. *prp2*$^{-/-}$ no treatment | 0.002367 | -0.02112 to 0.02585 | No | ns | >0.9999 | B-E |
| 16 | AB TBI vs. *prp2*$^{-/-}$ + TBI | -0.02586 | -0.04854 to -0.003180 | Yes | * | 0.0132 | B-F |
| 17 | AB TBI vs. *prp2*$^{-/-}$ 10uM MPEP | -0.01689 | -0.05295 to 0.01916 | No | ns | 0.8425 | B-G |
| 18 | AB TBI vs. *prp2*$^{-/-}$ 10uM MPEP + TBI | -0.1217 | -0.1546 to -0.08882 | Yes | **** | <0.0001 | B-H |
| 19 | AB 10uM MPEP vs. AB 10uM MPEP + TBI | -0.02341 | -0.06625 to 0.01944 | No | ns | 0.7078 | C-D |
| 20 | AB 10uM MPEP vs. *prp2*$^{-/-}$ no treatment | 0.007308 | -0.02663 to 0.04125 | No | ns | 0.9980 | C-E |
| 21 | AB 10uM MPEP vs. *prp2*$^{-/-}$ + TBI | -0.02092 | -0.05430 to 0.01247 | No | ns | 0.5431 | C-F |
| 22 | AB 10uM MPEP vs. *prp2*$^{-/-}$ 10uM MPEP | -0.01195 | -0.05555 to 0.03164 | No | ns | 0.9908 | C-G |
| 23 | AB 10uM MPEP vs. *prp2*$^{-/-}$ 10uM MPEP + TBI | -0.1168 | -0.1578 to -0.07576 | Yes | **** | <0.0001 | C-H |
| 24 | AB 10uM MPEP + TBI vs. *prp2*$^{-/-}$ no treatment | 0.03072 | -0.004776 to 0.06621 | No | ns | 0.1453 | D-E |
| 25 | AB 10uM MPEP + TBI vs. *prp2*$^{-/-}$ + TBI | 0.002491 | -0.03247 to 0.03746 | No | ns | >0.9999 | D-F |
| 26 | AB 10uM MPEP + TBI vs. *prp2*$^{-/-}$ 10uM MPEP | 0.01146 | -0.03336 to 0.05627 | No | ns | 0.9940 | D-G |
| 27 | AB 10uM MPEP + TBI vs. *prp2*$^{-/-}$ 10uM MPEP + TBI | -0.09336 | -0.1357 to -0.05105 | Yes | **** | <0.0001 | D-H |
| 28 | *prp2*$^{-/-}$ no treatment vs. *prp2*$^{-/-}$ + TBI | -0.02823 | -0.05144 to -0.005016 | Yes | ** | 0.0059 | E-F |
| 29 | *prp2*$^{-/-}$ no treatment vs. *prp2*$^{-/-}$ 10uM MPEP | -0.01926 | -0.05566 to 0.01713 | No | ns | 0.7404 | E-G |
| 30 | *prp2*$^{-/-}$ no treatment vs. *prp2*$^{-/-}$ 10uM MPEP + TBI | -0.1241 | -0.1573 to -0.09082 | Yes | **** | <0.0001 | E-H |
| 31 | *prp2*$^{-/-}$ + TBI vs. *prp2*$^{-/-}$ 10uM MPEP | 0.008964 | -0.02692 to 0.04484 | No | ns | 0.9948 | F-G |

### Analysis associated with Figure 6C

| | **Ordinary one-way ANOVA** Multiple comparisons | | | | | | | | |
|---|---|---|---|---|---|---|---|---|---|
| 32 | *prp2*$^{-/-}$ + TBI vs. *prp2*$^{-/-}$ 10uM MPEP + TBI | -0.09585 | -0.1286 to -0.06316 | Yes | **** | <0.0001 | F-H | | |
| 33 | *prp2*$^{-/-}$ 10uM MPEP vs. *prp2*$^{-/-}$ 10uM MPEP + TBI | -0.1048 | -0.1479 to -0.06175 | Yes | **** | <0.0001 | G-H | | |
| 34 | | | | | | | | | |
| 35 | **Test details** | **Mean 1** | **Mean 2** | **Mean Diff.** | **SE of diff.** | **n1** | **n2** | **q** | **DF** |
| 36 | AB no treatment vs. AB TBI | 0.04788 | 0.04474 | 0.003136 | 0.007620 | 56 | 59 | 0.5821 | 291 |
| 37 | AB no treatment vs. AB 10uM MPEP | 0.04788 | 0.04968 | -0.001805 | 0.01107 | 56 | 18 | 0.2306 | 291 |
| 38 | AB no treatment vs. AB 10uM MPEP + TBI | 0.04788 | 0.07309 | -0.02521 | 0.01158 | 56 | 16 | 3.080 | 291 |
| 39 | AB no treatment vs. *prp2*$^{-/-}$ no treatment | 0.04788 | 0.04237 | 0.005504 | 0.007790 | 56 | 54 | 0.9992 | 291 |
| 40 | AB no treatment vs. *prp2*$^{-/-}$ + TBI | 0.04788 | 0.07060 | -0.02272 | 0.007529 | 56 | 62 | 4.268 | 291 |
| 41 | AB no treatment vs. *prp2*$^{-/-}$ 10uM MPEP | 0.04788 | 0.06164 | -0.01376 | 0.01187 | 56 | 15 | 1.639 | 291 |
| 42 | AB no treatment vs. *prp2*$^{-/-}$ 10uM MPEP + TBI | 0.04788 | 0.1665 | -0.1186 | 0.01084 | 56 | 19 | 15.46 | 291 |
| 43 | AB TBI vs. AB 10uM MPEP | 0.04474 | 0.04968 | -0.004941 | 0.01100 | 59 | 18 | 0.6354 | 291 |
| 44 | AB TBI vs. AB 10uM MPEP + TBI | 0.04474 | 0.07309 | -0.02835 | 0.01151 | 59 | 16 | 3.483 | 291 |
| 45 | AB TBI vs. *prp2*$^{-/-}$ no treatment | 0.04474 | 0.04237 | 0.002367 | 0.007692 | 59 | 54 | 0.4353 | 291 |
| 46 | AB TBI vs. *prp2*$^{-/-}$ + TBI | 0.04474 | 0.07060 | -0.02586 | 0.007428 | 59 | 62 | 4.923 | 291 |
| 47 | AB TBI vs. *prp2*$^{-/-}$ 10uM MPEP | 0.04474 | 0.06164 | -0.01689 | 0.01181 | 59 | 15 | 2.023 | 291 |
| 48 | AB TBI vs. *prp2*$^{-/-}$ 10uM MPEP + TBI | 0.04474 | 0.1665 | -0.1217 | 0.01077 | 59 | 19 | 15.98 | 291 |
| 49 | AB 10uM MPEP vs. AB 10uM MPEP + TBI | 0.04968 | 0.07309 | -0.02341 | 0.01403 | 18 | 16 | 2.359 | 291 |
| 50 | AB 10uM MPEP vs. *prp2*$^{-/-}$ no treatment | 0.04968 | 0.04237 | 0.007308 | 0.01112 | 18 | 54 | 0.9298 | 291 |
| 51 | AB 10uM MPEP vs. *prp2*$^{-/-}$ + TBI | 0.04968 | 0.07060 | -0.02092 | 0.01094 | 18 | 62 | 2.705 | 291 |
| 52 | AB 10uM MPEP vs. *prp2*$^{-/-}$ 10uM MPEP | 0.04968 | 0.06164 | -0.01195 | 0.01428 | 18 | 15 | 1.184 | 291 |
| 53 | AB 10uM MPEP vs. *prp2*$^{-/-}$ 10uM MPEP + TBI | 0.04968 | 0.1665 | -0.1168 | 0.01343 | 18 | 19 | 12.29 | 291 |
| 54 | AB 10uM MPEP + TBI vs. *prp2*$^{-/-}$ no treatment | 0.07309 | 0.04237 | 0.03072 | 0.01163 | 16 | 54 | 3.737 | 291 |
| 55 | AB 10uM MPEP + TBI vs. *prp2*$^{-/-}$ + TBI | 0.07309 | 0.07060 | 0.002491 | 0.01145 | 16 | 62 | 0.3076 | 291 |
| 56 | AB 10uM MPEP + TBI vs. *prp2*$^{-/-}$ 10uM MPEP | 0.07309 | 0.06164 | 0.01146 | 0.01468 | 16 | 15 | 1.104 | 291 |
| 57 | AB 10uM MPEP + TBI vs. *prp2*$^{-/-}$ 10uM MPEP + TBI | 0.07309 | 0.1665 | -0.09336 | 0.01386 | 16 | 19 | 9.528 | 291 |
| 58 | *prp2*$^{-/-}$ no treatment vs. *prp2*$^{-/-}$ + TBI | 0.04237 | 0.07060 | -0.02823 | 0.007602 | 54 | 62 | 5.251 | 291 |
| 59 | *prp2*$^{-/-}$ no treatment vs. *prp2*$^{-/-}$ 10uM MPEP | 0.04237 | 0.06164 | -0.01926 | 0.01192 | 54 | 15 | 2.285 | 291 |
| 60 | *prp2*$^{-/-}$ no treatment vs. *prp2*$^{-/-}$ 10uM MPEP + TBI | 0.04237 | 0.1665 | -0.1241 | 0.01089 | 54 | 19 | 16.11 | 291 |
| 61 | *prp2*$^{-/-}$ + TBI vs. *prp2*$^{-/-}$ 10uM MPEP | 0.07060 | 0.06164 | 0.008964 | 0.01175 | 62 | 15 | 1.079 | 291 |
| 62 | *prp2*$^{-/-}$ + TBI vs. *prp2*$^{-/-}$ 10uM MPEP + TBI | 0.07060 | 0.1665 | -0.09585 | 0.01071 | 62 | 19 | 12.66 | 291 |

### Analysis associated with Figure 6C

| | **Ordinary one-way ANOVA** Multiple comparisons | | | | | | | | |
|---|---|---|---|---|---|---|---|---|---|
| 63 | *prp2*$^{-/-}$ 10uM MPEP vs. *prp2*$^{-/-}$ 10uM MPEP + TBI | 0.06164 | 0.1665 | -0.1048 | 0.01411 | 15 | 19 | 10.51 | 291 |
| 64 | | | | | | | | | |
| 65 | **Compact letter display** | | | | | | | | |
| 66 | *prp2*$^{-/-}$ 10uM MPEP + TBI | A | | | | | | | |
| 67 | AB 10uM MPEP + TBI | B C | | | | | | | |
| 68 | *prp2*$^{-/-}$ + TBI | B | | | | | | | |
| 69 | *prp2*$^{-/-}$ 10uM MPEP | B C | | | | | | | |
| 70 | AB 10uM MPEP | B C | | | | | | | |
| 71 | AB no treatment | B C | | | | | | | |
| 72 | AB TBI | C | | | | | | | |
| 73 | *prp2*$^{-/-}$ no treatment | C | | | | | | | |

### Analysis associated with Figure 6D

| | Ordinary one-way ANOVA<br>ANOVA results | | | | | |
|---|---|---|---|---|---|---|
| 1 | Table Analyzed | PC_TBI_AB_prp2_CHPG_5u | | | | |
| 2 | Data sets analyzed | A-H | | | | |
| 3 | | | | | | |
| 4 | **ANOVA summary** | | | | | |
| 5 | F | 4.421 | | | | |
| 6 | P value | 0.0001 | | | | |
| 7 | P value summary | *** | | | | |
| 8 | Significant diff. among means (P < 0.05)? | Yes | | | | |
| 9 | R squared | 0.09920 | | | | |
| 10 | | | | | | |
| 11 | **Brown-Forsythe test** | | | | | |
| 12 | F (DFn, DFd) | 4.228 (7, 281) | | | | |
| 13 | P value | 0.0002 | | | | |
| 14 | P value summary | *** | | | | |
| 15 | Are SDs significantly different (P < 0.05)? | Yes | | | | |
| 16 | | | | | | |
| 17 | **Bartlett's test** | | | | | |
| 18 | Bartlett's statistic (corrected) | 39.92 | | | | |
| 19 | P value | <0.0001 | | | | |
| 20 | P value summary | **** | | | | |
| 21 | Are SDs significantly different (P < 0.05)? | Yes | | | | |
| 22 | | | | | | |
| 23 | **ANOVA table** | **SS** | **DF** | **MS** | **F (DFn, DFd)** | **P value** |
| 24 | Treatment (between columns) | 0.03979 | 7 | 0.005685 | F (7, 281) = 4.421 | P=0.0001 |
| 25 | Residual (within columns) | 0.3613 | 281 | 0.001286 | | |
| 26 | Total | 0.4011 | 288 | | | |
| 27 | | | | | | |
| 28 | **Data summary** | | | | | |
| 29 | Number of treatments (columns) | 8 | | | | |
| 30 | Number of values (total) | 289 | | | | |

### Analysis associated with Figure 6D

| Ordinary one-way ANOVA Multiple comparisons | | | | | | | |
|---|---|---|---|---|---|---|---|
| 1 | Number of families | 1 | | | | | |
| 2 | Number of comparisons per family | 28 | | | | | |
| 3 | Alpha | 0.05 | | | | | |
| 4 | | | | | | | |
| 5 | **Tukey's multiple comparisons test** | **Mean Diff.** | **95.00% CI of diff.** | **Significant?** | **Summary** | **Adjusted P Value** | |
| 6 | AB no treatment vs. AB TBI | 0.003136 | -0.01730 to 0.02357 | No | ns | 0.9998 | A-B |
| 7 | AB no treatment vs. AB 5uM CHPG | 0.004899 | -0.02994 to 0.03974 | No | ns | 0.9999 | A-C |
| 8 | AB no treatment vs. AB 5uM CHPG + TBI | 0.01994 | -0.01766 to 0.05753 | No | ns | 0.7383 | A-D |
| 9 | AB no treatment vs. prp2-/-no treatment | 0.005504 | -0.01538 to 0.02639 | No | ns | 0.9928 | A-E |
| 10 | AB no treatment vs. prp2-/-+ TBI | -0.02272 | -0.04291 to -0.002533 | Yes | * | 0.0154 | A-F |
| 11 | AB no treatment vs. prp2-/- 5uM CHPG | 0.008346 | -0.02649 to 0.04318 | No | ns | 0.9960 | A-G |
| 12 | AB no treatment vs. prp2-/- 5uM CHPG + TBI | 0.006605 | -0.02011 to 0.03332 | No | ns | 0.9951 | A-H |
| 13 | AB TBI vs. AB 5uM CHPG | 0.001763 | -0.03292 to 0.03644 | No | ns | >0.9999 | B-C |
| 14 | AB TBI vs. AB 5uM CHPG + TBI | 0.01680 | -0.02065 to 0.05425 | No | ns | 0.8701 | B-D |
| 15 | AB TBI vs. prp2-/-no treatment | 0.002367 | -0.01826 to 0.02299 | No | ns | >0.9999 | B-E |
| 16 | AB TBI vs. prp2-/-+ TBI | -0.02586 | -0.04578 to -0.005941 | Yes | ** | 0.0023 | B-F |
| 17 | AB TBI vs. prp2-/- 5uM CHPG | 0.005209 | -0.02947 to 0.03989 | No | ns | 0.9998 | B-G |
| 18 | AB TBI vs. prp2-/- 5uM CHPG + TBI | 0.003469 | -0.02305 to 0.02998 | No | ns | >0.9999 | B-H |
| 19 | AB 5uM CHPG vs. AB 5uM CHPG + TBI | 0.01504 | -0.03185 to 0.06193 | No | ns | 0.9770 | C-D |
| 20 | AB 5uM CHPG vs. prp2-/-no treatment | 0.0006044 | -0.03435 to 0.03555 | No | ns | >0.9999 | C-E |
| 21 | AB 5uM CHPG vs. prp2-/-+ TBI | -0.02762 | -0.06216 to 0.006916 | No | ns | 0.2253 | C-F |
| 22 | AB 5uM CHPG vs. prp2-/- 5uM CHPG | 0.003446 | -0.04126 to 0.04816 | No | ns | >0.9999 | C-G |
| 23 | AB 5uM CHPG vs. prp2-/- 5uM CHPG + TBI | 0.001706 | -0.03701 to 0.04042 | No | ns | >0.9999 | C-H |
| 24 | AB 5uM CHPG + TBI vs. prp2-/-no treatment | -0.01443 | -0.05213 to 0.02327 | No | ns | 0.9400 | D-E |
| 25 | AB 5uM CHPG + TBI vs. prp2-/-+ TBI | -0.04266 | -0.07998 to -0.005340 | Yes | * | 0.0129 | D-F |
| 26 | AB 5uM CHPG + TBI vs. prp2-/- 5uM CHPG | -0.01159 | -0.05848 to 0.03530 | No | ns | 0.9951 | D-G |
| 27 | AB 5uM CHPG + TBI vs. prp2-/- 5uM CHPG + TBI | -0.01333 | -0.05455 to 0.02789 | No | ns | 0.9759 | D-H |
| 28 | prp2-/-no treatment vs. prp2-/-+ TBI | -0.02823 | -0.04861 to -0.007842 | Yes | *** | 0.0008 | E-F |
| 29 | prp2-/-no treatment vs. prp2-/- 5uM CHPG | 0.002842 | -0.03211 to 0.03779 | No | ns | >0.9999 | E-G |
| 30 | prp2-/-no treatment vs. prp2-/- 5uM CHPG + TBI | 0.001101 | -0.02577 to 0.02797 | No | ns | >0.9999 | E-H |
| 31 | prp2-/-+ TBI vs. prp2-/- 5uM CHPG | 0.03107 | -0.003470 to 0.06561 | No | ns | 0.1131 | F-G |

### Analysis associated with Figure 6D

| | Ordinary one-way ANOVA<br>Multiple comparisons | | | | | | | | |
|---|---|---|---|---|---|---|---|---|---|
| 32 | prp2-/-+ TBI vs. prp2-/- 5uM CHPG + TBI | 0.02933 | 0.003000 to 0.05566 | Yes | * | 0.0173 | F-H | | |
| 33 | prp2-/- 5uM CHPG vs. prp2-/- 5uM CHPG + TBI | -0.001741 | -0.04046 to 0.03698 | No | ns | >0.9999 | G-H | | |
| 34 | | | | | | | | | |
| 35 | **Test details** | **Mean 1** | **Mean 2** | **Mean Diff.** | **SE of diff.** | **n1** | **n2** | **q** | **DF** |
| 36 | AB no treatment vs. AB TBI | 0.04788 | 0.04474 | 0.003136 | 0.006690 | 56 | 59 | 0.6629 | 281 |
| 37 | AB no treatment vs. AB 5uM CHPG | 0.04788 | 0.04298 | 0.004899 | 0.01141 | 56 | 12 | 0.6074 | 281 |
| 38 | AB no treatment vs. AB 5uM CHPG + TBI | 0.04788 | 0.02794 | 0.01994 | 0.01231 | 56 | 10 | 2.290 | 281 |
| 39 | AB no treatment vs. prp2-/-no treatment | 0.04788 | 0.04237 | 0.005504 | 0.006839 | 56 | 54 | 1.138 | 281 |
| 40 | AB no treatment vs. prp2-/-+ TBI | 0.04788 | 0.07060 | -0.02272 | 0.006611 | 56 | 62 | 4.861 | 281 |
| 41 | AB no treatment vs. prp2-/- 5uM CHPG | 0.04788 | 0.03953 | 0.008346 | 0.01141 | 56 | 12 | 1.035 | 281 |
| 42 | AB no treatment vs. prp2-/- 5uM CHPG + TBI | 0.04788 | 0.04127 | 0.006605 | 0.008749 | 56 | 24 | 1.068 | 281 |
| 43 | AB TBI vs. AB 5uM CHPG | 0.04474 | 0.04298 | 0.001763 | 0.01136 | 59 | 12 | 0.2196 | 281 |
| 44 | AB TBI vs. AB 5uM CHPG + TBI | 0.04474 | 0.02794 | 0.01680 | 0.01226 | 59 | 10 | 1.937 | 281 |
| 45 | AB TBI vs. prp2-/-no treatment | 0.04474 | 0.04237 | 0.002367 | 0.006753 | 59 | 54 | 0.4958 | 281 |
| 46 | AB TBI vs. prp2-/-+ TBI | 0.04474 | 0.07060 | -0.02586 | 0.006522 | 59 | 62 | 5.607 | 281 |
| 47 | AB TBI vs. prp2-/- 5uM CHPG | 0.04474 | 0.03953 | 0.005209 | 0.01136 | 59 | 12 | 0.6488 | 281 |
| 48 | AB TBI vs. prp2-/- 5uM CHPG + TBI | 0.04474 | 0.04127 | 0.003469 | 0.008682 | 59 | 24 | 0.5650 | 281 |
| 49 | AB 5uM CHPG vs. AB 5uM CHPG + TBI | 0.04298 | 0.02794 | 0.01504 | 0.01535 | 12 | 10 | 1.385 | 281 |
| 50 | AB 5uM CHPG vs. prp2-/-no treatment | 0.04298 | 0.04237 | 0.0006044 | 0.01144 | 12 | 54 | 0.07469 | 281 |
| 51 | AB 5uM CHPG vs. prp2-/-+ TBI | 0.04298 | 0.07060 | -0.02762 | 0.01131 | 12 | 62 | 3.454 | 281 |
| 52 | AB 5uM CHPG vs. prp2-/- 5uM CHPG | 0.04298 | 0.03953 | 0.003446 | 0.01464 | 12 | 12 | 0.3329 | 281 |
| 53 | AB 5uM CHPG vs. prp2-/- 5uM CHPG + TBI | 0.04298 | 0.04127 | 0.001706 | 0.01268 | 12 | 24 | 0.1902 | 281 |
| 54 | AB 5uM CHPG + TBI vs. prp2-/-no treatment | 0.02794 | 0.04237 | -0.01443 | 0.01235 | 10 | 54 | 1.653 | 281 |
| 55 | AB 5uM CHPG + TBI vs. prp2-/-+ TBI | 0.02794 | 0.07060 | -0.04266 | 0.01222 | 10 | 62 | 4.937 | 281 |
| 56 | AB 5uM CHPG + TBI vs. prp2-/- 5uM CHPG | 0.02794 | 0.03953 | -0.01159 | 0.01535 | 10 | 12 | 1.068 | 281 |
| 57 | AB 5uM CHPG + TBI vs. prp2-/- 5uM CHPG + TBI | 0.02794 | 0.04127 | -0.01333 | 0.01350 | 10 | 24 | 1.397 | 281 |
| 58 | prp2-/-no treatment vs. prp2-/-+ TBI | 0.04237 | 0.07060 | -0.02823 | 0.006675 | 54 | 62 | 5.980 | 281 |
| 59 | prp2-/-no treatment vs. prp2-/- 5uM CHPG | 0.04237 | 0.03953 | 0.002842 | 0.01144 | 54 | 12 | 0.3512 | 281 |
| 60 | prp2-/-no treatment vs. prp2-/- 5uM CHPG + TBI | 0.04237 | 0.04127 | 0.001101 | 0.008797 | 54 | 24 | 0.1770 | 281 |
| 61 | prp2-/-+ TBI vs. prp2-/- 5uM CHPG | 0.07060 | 0.03953 | 0.03107 | 0.01131 | 62 | 12 | 3.885 | 281 |
| 62 | prp2-/-+ TBI vs. prp2-/- 5uM CHPG + TBI | 0.07060 | 0.04127 | 0.02933 | 0.008621 | 62 | 24 | 4.811 | 281 |

### Analysis associated with Figure 6D

| **Ordinary one-way ANOVA**<br>Multiple comparisons | | | | | | | | | |
|---|---|---|---|---|---|---|---|---|---|
| 63 | prp2-/- 5uM CHPG vs. prp2-/- 5uM CHPG + TBI | 0.03953 | 0.04127 | -0.001741 | 0.01268 | 12 | 24 | 0.1942 | 281 |

### Analysis associated with Supplementary Figure S1A

| | **Kruskal-Wallis test**<br>ANOVA results | |
|---|---|---|
| 1 | Table Analyzed | UA3171 AB42 long injected 3dpf |
| 2 | | |
| 3 | **Kruskal-Wallis test** | |
| 4 | P value | <0.0001 |
| 5 | Exact or approximate P value? | Approximate |
| 6 | P value summary | **** |
| 7 | Do the medians vary signif. (P < 0.05)? | Yes |
| 8 | Number of groups | 10 |
| 9 | Kruskal-Wallis statistic | 100.4 |
| 10 | | |
| 11 | **Data summary** | |
| 12 | Number of treatments (columns) | 10 |
| 13 | Number of values (total) | 372 |

### Analysis associated with Supplementary Figure S1A

| | **Kruskal-Wallis test**<br>Multiple comparisons | | | | | |
|---|---|---|---|---|---|---|
| 1 | Number of families | 1 | | | | |
| 2 | Number of comparisons per family | 45 | | | | |
| 3 | Alpha | 0.05 | | | | |
| 4 | | | | | | |
| 5 | **Dunn's multiple comparisons test** | **Mean rank diff.** | **Significant?** | **Summary** | **Adjusted P Value** | |
| 6 | Uninjected vs. Uninjected + 50uM MPEP | -0.2686 | No | ns | >0.9999 | A-B |
| 7 | Uninjected vs. Uninjected + 50uM CHPG | 13.19 | No | ns | >0.9999 | A-C |
| 8 | Uninjected vs. Mock inj | -40.79 | No | ns | >0.9999 | A-D |
| 9 | Uninjected vs. Mock injected + 50uM MPEP | -44.07 | No | ns | >0.9999 | A-E |
| 10 | Uninjected vs. Mock injected + 50uM CHPG | -30.95 | No | ns | >0.9999 | A-F |
| 11 | Uninjected vs. AB42 long inj | -118.5 | Yes | **** | <0.0001 | A-G |
| 12 | Uninjected vs. AB42 long inj + 50uM MPEP | -72.27 | Yes | ** | 0.0017 | A-H |
| 13 | Uninjected vs. AB42 long inj + 50uM CHPG | -107.3 | Yes | **** | <0.0001 | A-I |
| 14 | Uninjected vs. AB monomer inj | -4.674 | No | ns | >0.9999 | A-J |
| 15 | Uninjected + 50uM MPEP vs. Uninjected + 50uM CHPG | 13.46 | No | ns | >0.9999 | B-C |
| 16 | Uninjected + 50uM MPEP vs. Mock inj | -40.52 | No | ns | >0.9999 | B-D |
| 17 | Uninjected + 50uM MPEP vs. Mock injected + 50uM MPEP | -43.81 | No | ns | >0.9999 | B-E |
| 18 | Uninjected + 50uM MPEP vs. Mock injected + 50uM CHPG | -30.68 | No | ns | >0.9999 | B-F |
| 19 | Uninjected + 50uM MPEP vs. AB42 long inj | -118.2 | Yes | ** | 0.0011 | B-G |
| 20 | Uninjected + 50uM MPEP vs. AB42 long inj + 50uM MPEP | -72.00 | No | ns | 0.5827 | B-H |
| 21 | Uninjected + 50uM MPEP vs. AB42 long inj + 50uM CHPG | -107.0 | Yes | * | 0.0167 | B-I |
| 22 | Uninjected + 50uM MPEP vs. AB monomer inj | -4.405 | No | ns | >0.9999 | B-J |
| 23 | Uninjected + 50uM CHPG vs. Mock inj | -53.98 | No | ns | >0.9999 | C-D |
| 24 | Uninjected + 50uM CHPG vs. Mock injected + 50uM MPEP | -57.27 | No | ns | >0.9999 | C-E |
| 25 | Uninjected + 50uM CHPG vs. Mock injected + 50uM CHPG | -44.14 | No | ns | >0.9999 | C-F |
| 26 | Uninjected + 50uM CHPG vs. AB42 long inj | -131.7 | Yes | ** | 0.0013 | C-G |
| 27 | Uninjected + 50uM CHPG vs. AB42 long inj + 50uM MPEP | -85.46 | No | ns | 0.3709 | C-H |
| 28 | Uninjected + 50uM CHPG vs. AB42 long inj + 50uM CHPG | -120.5 | Yes | * | 0.0135 | C-I |
| 29 | Uninjected + 50uM CHPG vs. AB monomer inj | -17.87 | No | ns | >0.9999 | C-J |
| 30 | Mock inj vs. Mock injected + 50uM MPEP | -3.287 | No | ns | >0.9999 | D-E |
| 31 | Mock inj vs. Mock injected + 50uM CHPG | 9.837 | No | ns | >0.9999 | D-F |

### Analysis associated with Supplementary Figure S1A

| | **Kruskal-Wallis test** Multiple comparisons | | | | | | |
|---|---|---|---|---|---|---|---|
| 32 | Mock inj vs. AB42 long inj | -77.72 | Yes | * | 0.0112 | D-G | |
| 33 | Mock inj vs. AB42 long inj + 50uM MPEP | -31.48 | No | ns | >0.9999 | D-H | |
| 34 | Mock inj vs. AB42 long inj + 50uM CHPG | -66.48 | No | ns | 0.2452 | D-I | |
| 35 | Mock inj vs. AB monomer inj | 36.11 | No | ns | >0.9999 | D-J | |
| 36 | Mock injected + 50uM MPEP vs. Mock injected + 50uM CHPG | 13.12 | No | ns | >0.9999 | E-F | |
| 37 | Mock injected + 50uM MPEP vs. AB42 long inj | -74.43 | No | ns | 0.2063 | E-G | |
| 38 | Mock injected + 50uM MPEP vs. AB42 long inj + 50uM MPEP | -28.19 | No | ns | >0.9999 | E-H | |
| 39 | Mock injected + 50uM MPEP vs. AB42 long inj + 50uM CHPG | -63.20 | No | ns | >0.9999 | E-I | |
| 40 | Mock injected + 50uM MPEP vs. AB monomer inj | 39.40 | No | ns | >0.9999 | E-J | |
| 41 | Mock injected + 50uM CHPG vs. AB42 long inj | -87.56 | No | ns | 0.8152 | F-G | |
| 42 | Mock injected + 50uM CHPG vs. AB42 long inj + 50uM MPEP | -41.32 | No | ns | >0.9999 | F-H | |
| 43 | Mock injected + 50uM CHPG vs. AB42 long inj + 50uM CHPG | -76.32 | No | ns | >0.9999 | F-I | |
| 44 | Mock injected + 50uM CHPG vs. AB monomer inj | 26.28 | No | ns | >0.9999 | F-J | |
| 45 | AB42 long inj vs. AB42 long inj + 50uM MPEP | 46.24 | No | ns | 0.1302 | G-H | |
| 46 | AB42 long inj vs. AB42 long inj + 50uM CHPG | 11.23 | No | ns | >0.9999 | G-I | |
| 47 | AB42 long inj vs. AB monomer inj | 113.8 | Yes | **** | <0.0001 | G-J | |
| 48 | AB42 long inj + 50uM MPEP vs. AB42 long inj + 50uM CHPG | -35.01 | No | ns | >0.9999 | H-I | |
| 49 | AB42 long inj + 50uM MPEP vs. AB monomer inj | 67.59 | Yes | * | 0.0133 | H-J | |
| 50 | AB42 long inj + 50uM CHPG vs. AB monomer inj | 102.6 | Yes | **** | <0.0001 | I-J | |
| 51 | | | | | | | |
| 52 | **Test details** | **Mean rank 1** | **Mean rank 2** | **Mean rank diff.** | **n1** | **n2** | **Z** |
| 53 | Uninjected vs. Uninjected + 50uM MPEP | 126.2 | 126.5 | -0.2686 | 57 | 13 | 0.009227 |
| 54 | Uninjected vs. Uninjected + 50uM CHPG | 126.2 | 113.0 | 13.19 | 57 | 10 | 0.4063 |
| 55 | Uninjected vs. Mock inj | 126.2 | 167.0 | -40.79 | 57 | 25 | 1.795 |
| 56 | Uninjected vs. Mock injected + 50uM MPEP | 126.2 | 170.3 | -44.07 | 57 | 15 | 1.604 |
| 57 | Uninjected vs. Mock injected + 50uM CHPG | 126.2 | 157.1 | -30.95 | 57 | 7 | 0.8160 |
| 58 | Uninjected vs. AB42 long inj | 126.2 | 244.7 | -118.5 | 57 | 98 | 7.512 |
| 59 | Uninjected vs. AB42 long inj + 50uM MPEP | 126.2 | 198.5 | -72.27 | 57 | 60 | 4.126 |
| 60 | Uninjected vs. AB42 long inj + 50uM CHPG | 126.2 | 233.5 | -107.3 | 57 | 42 | 5.570 |
| 61 | Uninjected vs. AB monomer inj | 126.2 | 130.9 | -4.674 | 57 | 45 | 0.2475 |
| 62 | Uninjected + 50uM MPEP vs. Uninjected + 50uM CHPG | 126.5 | 113.0 | 13.46 | 13 | 10 | 0.3380 |

### Analysis associated with Supplementary Figure S1A

| Kruskal-Wallis test<br>Multiple comparisons | | | | | | | |
|---|---|---|---|---|---|---|---|
| 63 | Uninjected + 50uM MPEP vs. Mock inj | 126.5 | 167.0 | -40.52 | 13 | 25 | 1.251 |
| 64 | Uninjected + 50uM MPEP vs. Mock injected + 50uM MPEP | 126.5 | 170.3 | -43.81 | 13 | 15 | 1.221 |
| 65 | Uninjected + 50uM MPEP vs. Mock injected + 50uM CHPG | 126.5 | 157.1 | -30.68 | 13 | 7 | 0.6911 |
| 66 | Uninjected + 50uM MPEP vs. AB42 long inj | 126.5 | 244.7 | -118.2 | 13 | 98 | 4.230 |
| 67 | Uninjected + 50uM MPEP vs. AB42 long inj + 50uM MPEP | 126.5 | 198.5 | -72.00 | 13 | 60 | 2.485 |
| 68 | Uninjected + 50uM MPEP vs. AB42 long inj + 50uM CHPG | 126.5 | 233.5 | -107.0 | 13 | 42 | 3.560 |
| 69 | Uninjected + 50uM MPEP vs. AB monomer inj | 126.5 | 130.9 | -4.405 | 13 | 45 | 0.1477 |
| 70 | Uninjected + 50uM CHPG vs. Mock inj | 113.0 | 167.0 | -53.98 | 10 | 25 | 1.523 |
| 71 | Uninjected + 50uM CHPG vs. Mock injected + 50uM MPEP | 113.0 | 170.3 | -57.27 | 10 | 15 | 1.481 |
| 72 | Uninjected + 50uM CHPG vs. Mock injected + 50uM CHPG | 113.0 | 157.1 | -44.14 | 10 | 7 | 0.9459 |
| 73 | Uninjected + 50uM CHPG vs. AB42 long inj | 113.0 | 244.7 | -131.7 | 10 | 98 | 4.189 |
| 74 | Uninjected + 50uM CHPG vs. AB42 long inj + 50uM MPEP | 113.0 | 198.5 | -85.46 | 10 | 60 | 2.642 |
| 75 | Uninjected + 50uM CHPG vs. AB42 long inj + 50uM CHPG | 113.0 | 233.5 | -120.5 | 10 | 42 | 3.615 |
| 76 | Uninjected + 50uM CHPG vs. AB monomer inj | 113.0 | 130.9 | -17.87 | 10 | 45 | 0.5397 |
| 77 | Mock inj vs. Mock injected + 50uM MPEP | 167.0 | 170.3 | -3.287 | 25 | 15 | 0.1063 |
| 78 | Mock inj vs. Mock injected + 50uM CHPG | 167.0 | 157.1 | 9.837 | 25 | 7 | 0.2429 |
| 79 | Mock inj vs. AB42 long inj | 167.0 | 244.7 | -77.72 | 25 | 98 | 3.663 |
| 80 | Mock inj vs. AB42 long inj + 50uM MPEP | 167.0 | 198.5 | -31.48 | 25 | 60 | 1.396 |
| 81 | Mock inj vs. AB42 long inj + 50uM CHPG | 167.0 | 233.5 | -66.48 | 25 | 42 | 2.779 |
| 82 | Mock inj vs. AB monomer inj | 167.0 | 130.9 | 36.11 | 25 | 45 | 1.529 |
| 83 | Mock injected + 50uM MPEP vs. Mock injected + 50uM CHPG | 170.3 | 157.1 | 13.12 | 15 | 7 | 0.3028 |
| 84 | Mock injected + 50uM MPEP vs. AB42 long inj | 170.3 | 244.7 | -74.43 | 15 | 98 | 2.835 |
| 85 | Mock injected + 50uM MPEP vs. AB42 long inj + 50uM MPEP | 170.3 | 198.5 | -28.19 | 15 | 60 | 1.031 |
| 86 | Mock injected + 50uM MPEP vs. AB42 long inj + 50uM CHPG | 170.3 | 233.5 | -63.20 | 15 | 42 | 2.219 |
| 87 | Mock injected + 50uM MPEP vs. AB monomer inj | 170.3 | 130.9 | 39.40 | 15 | 45 | 1.395 |
| 88 | Mock injected + 50uM CHPG vs. AB42 long inj | 157.1 | 244.7 | -87.56 | 7 | 98 | 2.363 |
| 89 | Mock injected + 50uM CHPG vs. AB42 long inj + 50uM MPEP | 157.1 | 198.5 | -41.32 | 7 | 60 | 1.092 |
| 90 | Mock injected + 50uM CHPG vs. AB42 long inj + 50uM CHPG | 157.1 | 233.5 | -76.32 | 7 | 42 | 1.974 |
| 91 | Mock injected + 50uM CHPG vs. AB monomer inj | 157.1 | 130.9 | 26.28 | 7 | 45 | 0.6829 |
| 92 | AB42 long inj vs. AB42 long inj + 50uM MPEP | 244.7 | 198.5 | 46.24 | 98 | 60 | 2.979 |
| 93 | AB42 long inj vs. AB42 long inj + 50uM CHPG | 244.7 | 233.5 | 11.23 | 98 | 42 | 0.6433 |

### Analysis associated with Supplementary Figure S1A

| Kruskal-Wallis test<br>Multiple comparisons | | | | | | | |
|---|---|---|---|---|---|---|---|
| 94 | AB42 long inj vs. AB monomer inj | 244.7 | 130.9 | 113.8 | 98 | 45 | 6.675 |
| 95 | AB42 long inj + 50uM MPEP vs. AB42 long inj + 50uM CHPG | 198.5 | 233.5 | -35.01 | 60 | 42 | 1.837 |
| 96 | AB42 long inj + 50uM MPEP vs. AB monomer inj | 198.5 | 130.9 | 67.59 | 60 | 45 | 3.619 |
| 97 | AB42 long inj + 50uM CHPG vs. AB monomer inj | 233.5 | 130.9 | 102.6 | 42 | 45 | 5.050 |
| 98 | | | | | | | |
| 99 | **Compact letter display** | | | | | | |
| 100 | AB42 long inj | A | | | | | |
| 101 | AB42 long inj + 50uM CHPG | A B | | | | | |
| 102 | AB42 long inj + 50uM MPEP | A B C | | | | | |
| 103 | Mock inj | B D | | | | | |
| 104 | Mock injected + 50uM MPEP | A D | | | | | |
| 105 | Mock injected + 50uM CHPG | A D | | | | | |
| 106 | Uninjected + 50uM MPEP | C D | | | | | |
| 107 | AB monomer inj | D | | | | | |
| 108 | Uninjected | D | | | | | |
| 109 | Uninjected + 50uM CHPG | C D | | | | | |

### Analysis associated with Supplementary Figure S1B

| | **Kruskal-Wallis test**<br>ANOVA results | |
|---|---|---|
| 1 | Table Analyzed | AB42 long + SAM (23-25hrs treat) |
| 2 | | |
| 3 | **Kruskal-Wallis test** | |
| 4 | P value | <0.0001 |
| 5 | Exact or approximate P value? | Approximate |
| 6 | P value summary | **** |
| 7 | Do the medians vary signif. (P < 0.05)? | Yes |
| 8 | Number of groups | 5 |
| 9 | Kruskal-Wallis statistic | 24.02 |
| 10 | | |
| 11 | **Data summary** | |
| 12 | Number of treatments (columns) | 5 |
| 13 | Number of values (total) | 181 |

### Analysis associated with Supplementary Figure S1B

| | **Kruskal-Wallis test** Multiple comparisons | | | | | | |
|---|---|---|---|---|---|---|---|
| 1 | Number of families | 1 | | | | | |
| 2 | Number of comparisons per family | 10 | | | | | |
| 3 | Alpha | 0.05 | | | | | |
| 4 | | | | | | | |
| 5 | **Dunn's multiple comparisons test** | **Mean rank diff.** | **Significant?** | **Summary** | **Adjusted P Value** | | |
| 6 | uninjected vs. AB long injected | -41.58 | Yes | **** | <0.0001 | A-B | |
| 7 | uninjected vs. ABL + 0.25uM SAM | -19.51 | No | ns | 0.5372 | A-C | |
| 8 | uninjected vs. ABL + 1uM SAM | -15.63 | No | ns | 0.8170 | A-D | |
| 9 | uninjected vs. ABL + 2.5uM SAM | -5.977 | No | ns | >0.9999 | A-E | |
| 10 | AB long injected vs. ABL + 0.25uM SAM | 22.06 | No | ns | 0.3450 | B-C | |
| 11 | AB long injected vs. ABL + 1uM SAM | 25.95 | No | ns | 0.0545 | B-D | |
| 12 | AB long injected vs. ABL + 2.5uM SAM | 35.60 | Yes | * | 0.0197 | B-E | |
| 13 | ABL + 0.25uM SAM vs. ABL + 1uM SAM | 3.886 | No | ns | >0.9999 | C-D | |
| 14 | ABL + 0.25uM SAM vs. ABL + 2.5uM SAM | 13.54 | No | ns | >0.9999 | C-E | |
| 15 | ABL + 1uM SAM vs. ABL + 2.5uM SAM | 9.650 | No | ns | >0.9999 | D-E | |
| 16 | | | | | | | |
| 17 | **Test details** | **Mean rank 1** | **Mean rank 2** | **Mean rank diff.** | **n1** | **n2** | **Z** |
| 18 | uninjected vs. AB long injected | 74.10 | 115.7 | -41.58 | 51 | 43 | 4.725 |
| 19 | uninjected vs. ABL + 0.25uM SAM | 74.10 | 93.61 | -19.51 | 51 | 27 | 1.929 |
| 20 | uninjected vs. ABL + 1uM SAM | 74.10 | 89.73 | -15.63 | 51 | 40 | 1.741 |
| 21 | uninjected vs. ABL + 2.5uM SAM | 74.10 | 80.08 | -5.977 | 51 | 20 | 0.5330 |
| 22 | AB long injected vs. ABL + 0.25uM SAM | 115.7 | 93.61 | 22.06 | 43 | 27 | 2.114 |
| 23 | AB long injected vs. ABL + 1uM SAM | 115.7 | 89.73 | 25.95 | 43 | 40 | 2.779 |
| 24 | AB long injected vs. ABL + 2.5uM SAM | 115.7 | 80.08 | 35.60 | 43 | 20 | 3.095 |
| 25 | ABL + 0.25uM SAM vs. ABL + 1uM SAM | 93.61 | 89.73 | 3.886 | 27 | 40 | 0.3671 |
| 26 | ABL + 0.25uM SAM vs. ABL + 2.5uM SAM | 93.61 | 80.08 | 13.54 | 27 | 20 | 1.080 |
| 27 | ABL + 1uM SAM vs. ABL + 2.5uM SAM | 89.73 | 80.08 | 9.650 | 40 | 20 | 0.8291 |
| 28 | | | | | | | |
| 29 | **Compact letter display** | | | | | | |
| 30 | AB long injected | A | | | | | |
| 31 | ABL + 0.25uM SAM | A B | | | | | |

### Analysis associated with Supplementary Figure S1B

| **Kruskal-Wallis test**<br>Multiple comparisons | | |
|---|---|---|
| 32 | ABL + 1uM SAM | A B |
| 33 | ABL + 2.5uM SAM | B |
| 34 | uninjected | B |
